# Tempo of gene regulation in wild and cultivated *Vitis* species shows coordination between cold deacclimation and budbreak

**DOI:** 10.1101/528828

**Authors:** Alisson P. Kovaleski, Jason P. Londo

## Abstract

**Highlight:** Faster deacclimation and budbreak phenology is related to a faster regulon rather than higher expression of specific genes. ABA is a master regulator of deacclimation.

**Abstract:** Dormancy release, loss of cold hardiness and budbreak are critical aspects of the annual cycle of deciduous perennial plants. Molecular control of these processes is not fully understood, and genotypic variation may be important for climate adaptation. Single-node cuttings from wild (*Vitis amurensis, V. riparia*) and cultivated *Vitis* genotypes (*V. vinifera* ‘Cabernet Sauvignon’, ‘Riesling’) were collected from the field during winter and placed under forcing conditions. Cold hardiness was measured daily, and buds were collected for RNA-Seq until budbreak. Field-collected single-node cuttings of ‘Riesling’ were treated with abscisic acid (ABA), and cold hardiness and budbreak at 7 °C were tracked. Wild *Vitis* genotypes had faster deacclimation and budbreak than *V. vinifera*. Temperature-sensing related genes were quickly and synchronously differentially expressed in all genotypes. ABA synthesis was down-regulated in all genotypes, and exogenous ABA prevented deacclimation. Ethylene- and oxidative stress-related genes were transiently up-regulated. Growth-related genes were up-regulated and showed staggering similar to deacclimation and budbreak of the four genotypes. The gene expression cascade that occurs during deacclimation and budburst phenology of fast (wild) and slow (cultivated) grapevines appears coordinated and temporally conserved. This may extend to other temperate woody species and suggest constraints on identification of process-specific keystone genes.

## Introduction

Low temperatures during winter are the most important factor limiting plant distribution (Burke *et al*., 1976). Perennial plants withstand low temperatures by forming wood and overwintering structures (i.e., dormant buds), utilizing dormancy to survive during non-growth-conducive environmental conditions. Dormancy onset is induced by changing day length, or day length and low temperatures (Cooke *et al*., 2012), and the transition from endogenous growth suppression (endodormancy) to environmental growth suppression (ecodormancy) occurs through accumulation of time in low, non-freezing temperatures – termed chilling accumulation (Lang *et al*., 1987; Cooke *et al*., 2012). Chilling greatly impacts timing of budbreak, and therefore budbreak phenology depends on temperatures during both winter (chilling) and spring (growing degree-days). This temperature-controlled interplay has been extensively studied in plants (Prentice *et al*., 1992). While dormant buds have been the subject of many studies regarding dormancy control (Olukolu *et al*., 2009; Ophir *et al*., 2009; van Dyk *et al*., 2010; Hedley *et al*., 2010; Díaz-Riquelme *et al*., 2012; Busov *et al*., 2016; Sudawan *et al*., 2016; Wu *et al*., 2017; Meitha *et al*., 2018), these have not considered the potential role of deacclimation, the loss of winter cold hardiness, as it impacts budbreak. Despite the foundation of dormancy research and importance of this trait for changing climate, the majority of cold hardiness studies focus on chill or freeze stress in annuals or actively growing tissues of perennials (Ruelland *et al*., 2002; Li *et al*., 2004; Nguyen *et al*., 2017; Liu *et al*., 2017a, b; Ban *et al*., 2017; Londo *et al*., 2018).

Dormancy is required for development of deep cold hardiness, and many processes associated with abiotic stress may be critical during dormancy. Transcriptomic tools (e.g., RNA-Seq) have revealed some of the gene expression processes that occur in dormant tissues. Dehydration-responsive element-binding 1 (DREB1)/C-repeat-binding factors (CBF) are known elicitors of cold-responsive genes following low temperature perception (Liu *et al*., 1998). Other stress related transcription factors [other DREBs, heat shock factors (HSFs), NAC and WRKY families] are also involved in the signaling, much of these shared between heat, drought, and biotic stresses (Liu *et al*., 1998, 2017a; Birkenbihl *et al*., 2012). The circadian clock is also likely involved in regulation of bud dormancy (Penfield, 2008; Cooke *et al*., 2012), and dormancy release and budbreak have shared processes with germination, such as hypoxia signaling via ETHYLENE-RESPONSIVE TRANSCRIPTION FACTORS (ERFs) regulation (Penfield, 2008; Meitha *et al*., 2018). Although known as central in senescence and dormancy, the role of abscisic acid (ABA) in bud dormancy is not fully understood (Cooke *et al*., 2012). ABA promotes the induction of dormancy by blocking cell-cell communication (Tylewicz *et al*., 2018). Exogenous ABA applications during the growing season have been shown to enhance hardiness earlier in the fall (Li and Dami, 2016) through increased expression of DREB1/CBF TFs (Rubio *et al*., 2018), and budbreak can be delayed by ABA when applied in dormant buds (Zheng *et al*., 2015). Considering recent evidence that budbreak phenology follows similar kinetics as deacclimation (Kovaleski *et al*., 2018), elucidating key gene regulation cascades during these processes could help identify candidate genes for climate adaptation.

Cold hardiness kinetics and budbreak phenology are tightly linked (Kovaleski *et al*., 2018), and understanding these two related traits may contribute to increased sustainability of crop species as climate variation increases, especially when acute cold weather episodes are expected to increase (Kolstad *et al*., 2010). As buds transition between dormancy stages, they alter the metabolic availability of sugars, carbon, and reactive oxygen species (ROS) without any visual cues (Lang *et al*., 1987). Therefore, understanding the regulation and timing of these changes requires an understanding of gene expression cascades. One major limitation to understanding important gene expression changes during dormancy arises from the time scale of dormancy, spanning throughout the entire winter. Traditional pairwise contrasts may be problematic over large time scales as subtle changes in expression will be undetectable between sample points, but significant over time. However, pairwise contrasts continue to be the routine method for determining differentially expressed genes in time-series studies (Fu *et al*., 2018; Meitha *et al*., 2018). New methods of examining gene expression data are needed to detect those changes that are significant over long periods of time.

Grapevine is the highest production value fruit crop worldwide with an estimated 67B (USD) impact (FAO, 2018). Sustainability is expected to be severely challenged by climate change even in traditional production regions (Cook and Wolkovich, 2016; Leolini *et al*., 2018). Phenology modelling of cultivated grapevine, *Vitis vinifera*, demonstrates wide variation in budbreak timing (Leolini *et al*., 2018) but genetic control of phenology is unknown in grapevine. Wild grapevine species that are frequently used in breeding programs to increase cold hardiness have higher rates of deacclimation during the spring when compared to *V. vinifera* (Londo and Kovaleski, 2017; Kovaleski *et al*., 2018). Comparisons between *Vitis* species could help identify molecular drivers of cold hardiness and deacclimation. In addition, the quantitative nature of deacclimation also responds to chill accumulation (Kovaleski *et al*., 2018). Therefore, understanding the genetic regulation of cold hardiness loss could elucidate some of the mechanisms that control dormancy in buds.

This study leverages phenological differences between species (*V. amurensis, V. riparia, V. vinifera*) and between cultivars within species (*V. vinifera* ‘Cabernet Sauvignon’ and ‘Riesling’) to uncover key aspects of gene regulation that occurs during deacclimation. The objective was to determine the transcriptional regulation and relationship between loss of cold hardiness and growth. We hypothesized that genotypes with early spring budbreak phenology may have faster transcriptomic responses tied to growth and development when placed under forcing conditions when contrasted with slower phenology genotypes. Alternatively, differences between genotypes with rapid or slow phenology could be due to acceleration of gene regulation cascades, or acceleration of “bottleneck” genes which trigger specific regulatory cascades. Additionally, we hypothesize that genes related to maintaining the cold hardy and dormant phenotype will be quickly down-regulated in all genotypes; genes related to deacclimation will either be transiently expressed or have a slow increase or decrease, but mostly synchronized between genotypes; and genes related to growth and budbreak will be staggered. Finally, we explore the use of ABA as a deacclimation inhibitor.

## Materials and Methods

### Plant material

Canes of four genotypes of grapevine were used in this study: two wild species, *V. amurensis* (PI588635) and *V. riparia* (PI588275) were collected from the USDA Plant Genetic Resources Unit in Geneva, New York; and two cultivars of cultivated grapevines (*V. vinifera*), ‘Cabernet Sauvignon’ and ‘Riesling’, from a local vineyard (42.845N, 77.004W). The two locations are approximately 6 km apart. The three species represent major clades in *Vitis:* American, Eurasian, and Asian (Fig. 1A). The *V. riparia* accession was originally collected in North Dakota, USA (https://www.arsgrin.gov/), while *V. vinifera* ‘Cabernet Sauvignon’ and ‘Riesling’ are cultivars domesticated in France and Germany, respectively. *V. amurensis* is native to temperate climates in East Asia, although no collection location information was available. Dormant canes were collected in mid-winter [February 2015; 1030 chill units (North Carolina model; Shaltout and Unrath, 1983)], and chopped into single node cuttings. The cuttings were placed with cut ends in cups of water and placed within a growth room with forcing conditions: 22 °C and 16/8h light/dark.

**Fig. 1.**
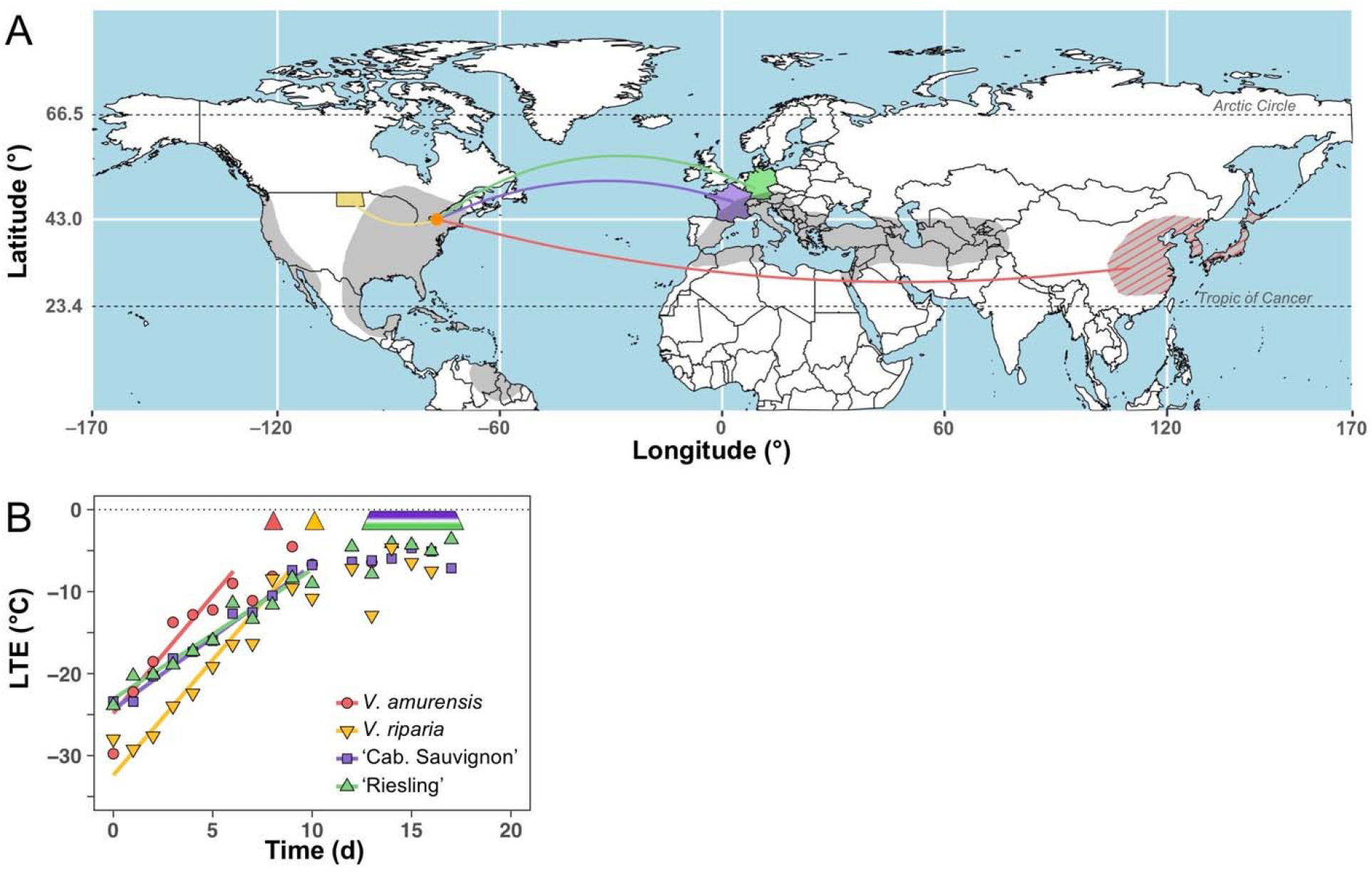
Distribution, cold hardiness and deacclimation of four *Vitis* spp. genotypes. (a) Distribution map of the centers of diversity for *Vitis* spp. in gray – American, Eurasian, and Asian (Redrawn from Alleweldt *et al*., 1990; Wan *et al*., 2013). Colored area denotes origin for each genotype used: *V. vinifera* ‘Cabernet Sauvignon’ and ‘Riesling’ were domesticated in France and Germany, respectively; *V. amurensis* (PI588635) is native to temperate climates in East Asia; and *V. riparia* (PI588275) was originally collected from the wild in North Dakota, USA. Orange dot shows where genotypes are cultivated and were collected from (43N, 77W). (b) Cold hardiness, indicated by symbols and predicted linear deacclimation for each genotype. Arrow heads (*V. amurensis* and *V. riparia*) or trapezoid (*V. vinifera*) at 0 °C line indicate timing of budbreak (E-L stage 3; Coombe and Iland, 2005). Deacclimation rates were 2.87, 2.81, 1.76, and 1.58 °C day^−1^ for *V. amurensis*, *V. riparia*, and *V. vinifera* ‘Cabernet Sauvignon’ and ‘Riesling’, respectively.

### Loss of cold hardiness and budbreak

Measurement of bud cold hardiness was conducted following standard differential thermal analysis (DTA) methods to detect lethal freezing temperature of grapevine buds (Mills *et al*., 2006). Buds were excised from canes, placed on thermoelectric modules and subjected to decreasing temperatures at −4 °C hour^−1^ (n=8). The release of heat that results from freezing of water inside the bud, the low temperature exotherm (LTE), was recorded using a Keithley data logger (Tektronix, Beaverton, OR). Cold hardiness was assessed over several days and rates were determined using linear regression of LTE data. A subsample of 10 replicate single node cuttings were left to fully deacclimate and reach budbreak. Phenology was visually assessed, and budbreak was noted when buds reached stage 3 of the modified E-L scale (Coombe and Iland, 2005).

### RNAseq Libraries and data analysis

Three replicates of three buds each were collected directly into LN, daily for each genotype, until buds reached E-L stage 3; corresponding to day 10, 12, 15, and 17 for *V. amurensis, V. riparia*, ‘Riesling’, and ‘Cabernet Sauvignon’, respectively. Day 0 samples were field-collected. Total RNA was extracted using Sigma Spectrum kits (Sigma-Aldrich, St. Louis, MO, USA), and strand-specific libraries were prepared (Borodina *et al*., 2011). Libraries were multiplexed (25-plex) and sequenced at 100 bp single-end reads on HiSeq 2500 (Illumina, Inc., San Diego, CA, USA) at Cornell University’s Institute of Biotechnology (Ithaca, NY, USA).

Raw reads were de-multiplexed using FastX_BarcodeSplitter.pl (Gordon and Hannon, 2010), quality trimmed using Cutadapt (Martin, 2011), aligned to the *V. vinifera* 12xV2 genome (https://urgi.versailles.inra.fr/Species/Vitis) and the V3 annotation (Canaguier *et al*., 2017) using STAR (Dobin *et al*., 2013), and uniquely aligned reads were quantified using HTSeq-count (Anders *et al*., 2015). Sequence data are available at NCBI-GEO (https://www.ncbi.nlm.nih.gov/geo/), series entry GSE124820.

Within each genotype, counts were normalized and grouped by days using the *edgeR* library in R (ver. 3.3.0, R Foundation for Statistical Computing). Multidimensional scaling and correlation plots were produced to visually verify outliers in libraries. After outlier removal, remaining data were analyzed using the *DESeq2* library (Love *et al*., 2014). In *DESeq2*, a full model containing a fourth order polynomial for time (t) as a continuous variable and intercept, was compared to a null model of intercept only, using likelihood ratio test. Genes with an adjusted-*P*≤0.10 [FDR correction (Benjamini and Hochberg, 1995)] were used for subsequent filtering. Using parameter estimates extracted from *DESeq2*, predicted normalized counts [counts per million (CPM)] were calculated for the time span of each genotype’s experiment (e.g., 0-10 days for *V. amurensis*, 0-17 days for *V. vinifera* ‘Cabernet Sauvignon’, as integers). Considering that counts for any given gene are a function of time (e.g., *CPM(t)*= *β_0_* + *β_1_t* + *β_2_t^2^* + *β_3_t^3^* + *β_4_t^4^*), the filters used were: *max{CPM(t)* | *0≤t≤t_f_}≥10* (i.e., maximum predicted CPM at any time point ≥10), where t_f_ is the final time point for any given genotype (e.g., for ‘Cabernet Sauvignon’, t_f_=10); and *log_2_*(*max*{*CPM*(*t*) | *0≤ t ≤ t_f_*}/*min*{*CPM*(*t*) | *0 ≤ t ≤ t_f_*})≥2 (i.e., logFC≥2). Remaining genes were considered relevant differentially expressed genes (DEGs).

Predicted logFC for each day calculated using *DESeq2* model estimates were used for cluster analysis of DEGs using Cluster 3.0 (ver. 3.0, Human Genome Center, University of Tokyo). Data were centered based on gene means, then on array means, followed by normalization by genes and arrays. Twenty *k*-Means clusters were used for genes, using Euclidian distances as the similarity metric, and 100 runs. The average behavior of the genes placed in each cluster was visually inspected, and placed into 4 general categories: *up-regulated*, relative expression increasing over time; *down-regulated*, relative expression decreasing over time; *transient* expression, peak expression occurred within the duration of experiment; and *random*, where most of the clusters had genes with a transient low expression within the duration of the experiment.

The resultant DEGs for each genotype were tested for pathway enrichment using VitisPathways (Osier, 2016), which leverages biological pathways and molecular functions in grapevine from the VitisNet database (Grimplet *et al*., 2009). Enriched pathways were determined using 1000 permutations, and a permuted-*P* <0.10. Pathway enrichment analysis was conducted for each genotype using all DEGs, as well as DEGs in the up-regulated, down-regulated, and transient expression groups. Enriched pathways that were shared between genotypes were observed using Cytoscape (ver. 3.6.1; Shannon *et al*., 2003). To plot trends in gene expression for pathways of interest, genes predicted to encode or encoding the same protein were aggregated, and mean expression was used for each day. Therefore, protein names are used as the notation to refer to expression levels, with few exceptions (e.g., transcription factors). Qualitative comparisons of expression behavior were done on a relative basis, using the maximum mean counts within the duration of the experiment as 100% for each genotype.

### ABA treatment of single node cuttings

Single node cuttings of *V. vinifera* ‘Riesling’ were prepared from material collected from the same vineyard in 4 April 2017 [1630 chill units (Shaltout and Unrath, 1983)]. The cuttings were placed in a growth chamber with average temperature of 7 °C (2–7–12–7 °C for 6h each) and a 0/24h light/dark photoperiod. ABA [ProTone^®^ SG (20% S-ABA), Valent BioSciences Corp., Libertyville, USA] treatments were applied as the solution inside cups where cuttings were placed. Two concentrations were used, 1mM and 5mM, and a distilled water control. After 42 days, remaining cuttings were washed in running water, the bottom ~1 cm of stems was cut, and cuttings were placed in cups with water and moved to forcing conditions (22 °C, 16h/8h light/dark). Cold hardiness was measured twice a week until 38 days, and three more sample dates of unequal distance until day 56 (or two for the control). Phenology was evaluated in 5 buds for each treatment using the modified E-L scale (Coombe and Iland, 2005) until buds reached stage 8 (2 separated leaves). Pairwise treatment comparisons were evaluated for both cold hardiness (day 0 through 38) and phenology (day 27 through 56) within days and means were separated using Tukey’s HSD test (α=0.01). A logistic regression was used to compare when each treatment reached budbreak (E-L stage 3), using the *drc* library (Ritz *et al*., 2015).

## Results

### Deacclimation and budbreak (RNA-Seq)

Wild species had a higher initial cold hardiness upon collection (−29.8 and −28.0 °C for *V. amurensis* and *V. riparia*, respectively) compared to *V. vinifera* genotypes (−23.4 and −23.9 °C for ‘Cabernet Sauvignon’ and ‘Riesling’, respectively; Fig. 1B). Both wild species also had higher deacclimation rates than the two *V. vinifera* genotypes (2.87, 2.81, 1.76, and 1.58 °C day^−1^ for *V. amurensis, V. riparia*, and *V. vinifera* ‘Cab. Sauvignon’ and ‘Riesling’, respectively). Budbreak occurred latest and sparsely on *V. vinifera* genotypes, starting at 13 days and spanning through day 17 for ‘Riesling’ and ‘Cabernet Sauvignon’ (Fig. 1B). Budbreak under forcing conditions occurred after 8 days in *V. amurensis* and 10 days for *V. riparia*.

### Analyses of Differential Gene Expression

The number of DEGs ranged from 8326 (*V. riparia*) to 9123 (*V. vinifera* ‘Cabernet Sauvignon’; Supplementary Fig. S1A). DEGs uniquely expressed in each species were 1348 (16.0%) for *V. amurensis*, 1006 (12%) for *V. riparia*, and 2232 (21.7%) for *V. vinifera* (‘Cabernet Sauvignon’ + ‘Riesling’). The majority of DEGs, 4551, were shared among all four genotypes. Cluster analysis of shared DEGS grouped 1192 DEGs as down-regulated, and 1065 DEGs as up-regulated (Supplementary Fig. S1B, C). Principal components (PC) 1 and 2 generated through prcomp() [within pcaPlot() using ntop=1000] separate global gene expression data points explaining 61% and 11% of the variance, respectively, when using all genotypes. Plotting PC2 clearly separates *V. vinifera* genotypes from *V. amurensis* and *V. riparia* (Fig. 2A, B). A third PC (*not shown;* 6% variance) separates *V. amurensis* from *V. riparia*. There appears to be a discontinuity at the higher end of PC1 when points are compared by time, where the variance is more closely related between the final timepoints in *V. vinifera* (15 d and 17 d for ‘Riesling’ and ‘Cabernet Sauvignon’, respectively) and the final timepoints in *V. amurensis* (10 d) and *V. riparia* (12 d) (Fig. 2A). Time was transformed to a thermal time measurement using deacclimation rate to normalize the response of each genotype to temperature, therefore creating a thermal time unit of *Deacclimation Degree-Days* [*DDD* = *k*_deacc_ × *time (d)*], where *k*_deacc_ is the rate of deacclimation. Timepoint transformation to DDD better scales the differences between genotypes in the distribution of PC1 (Fig. 2B). Global trends in gene expression are also similar across genotypes in respect to phenology as observed in heatmap of gene expression (Supplementary Fig. **S2**). A simple linear regression of PC1 by DDD results in an r^2^=78.4% (Fig. 2C), further demonstrating that the majority of the variation in gene expression comes from a time component, and that differences between the genotypes are derived from their inherent faster (wild species) or slower (cultivated species) response to temperature as measured by the deacclimation rate.

**Fig. 2.**
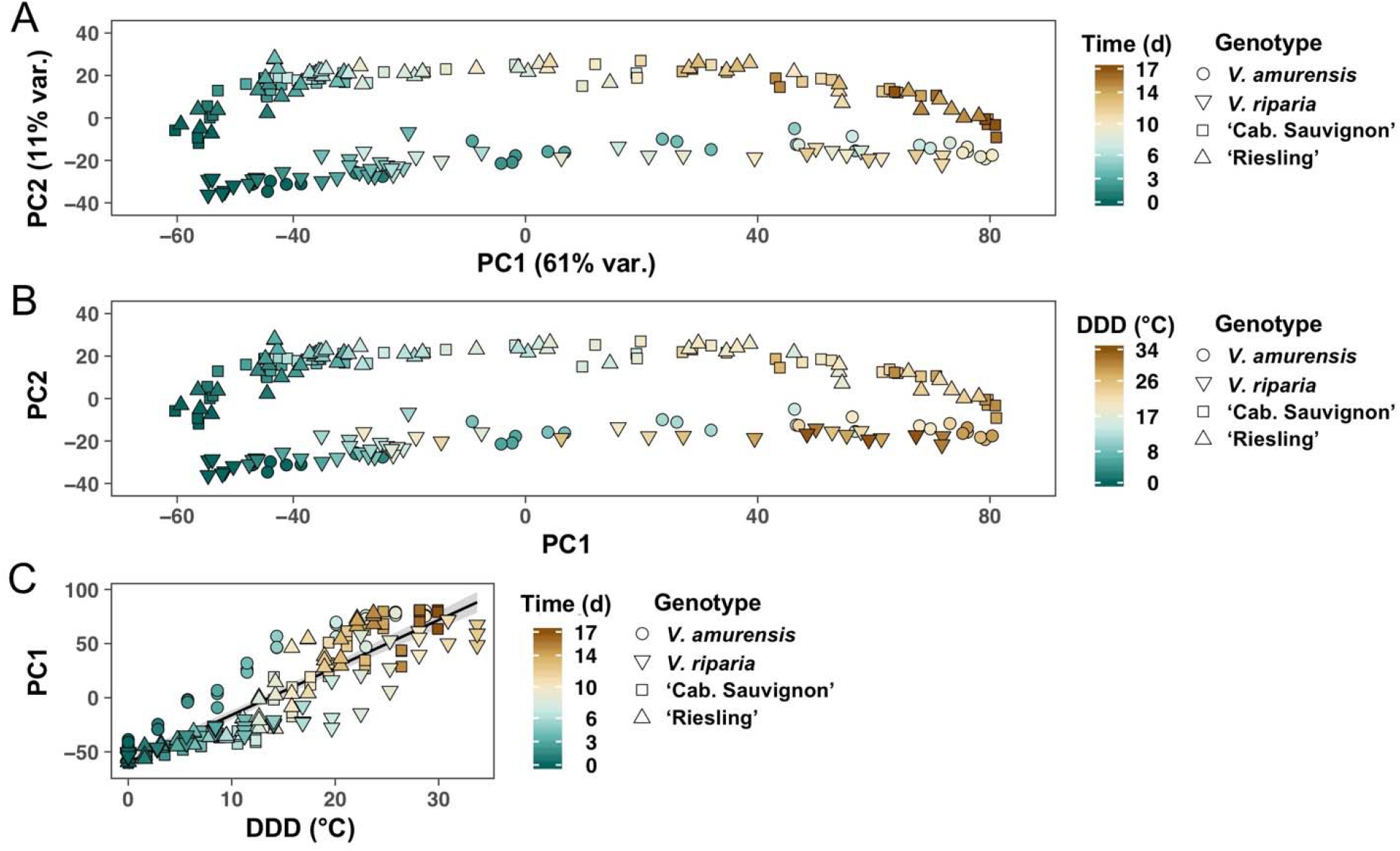
Principal components (PCs) for differential expression of four *Vitis* spp. genotypes. Principal components were generated using prcomp() and variance stabilized normalized counts. Distribution of genotypes and data points in PC1 and PC2, with points colored by (a) time in days and (b) thermal time in deacclimation degree-days [*DDD* = *k*_deacc_ × *time*, where *k*_deacc_ is the rate of deacclimation]. (c) Linear association of PC1 and DDD (r^2^ = 78.4%).

Pathway enrichment analysis was performed with gene lists composed of all DEGs to examine pathways where both up and down regulation may be occurring, but also for those in the down-regulated, up-regulated, and transiently expressed DEGs groups. There were 21 enriched pathways when all genes were queried, 22 for down-regulated patterns, 15 for up-regulated patterns, and 5 for transient patterns that were shared among all genotypes (Table 1). Overall, there were 43 enriched pathways that were shared among all genotypes: 16 were defined as associated with Metabolism, four with Environmental Information Processing, two with Cellular Processes, four with Transport, and 17 with Transcription Factors. Important pathways identified included many related to plant hormone production and signaling, sugar metabolism, and growth-related processes like cell cycling and photosynthesis. Critical genes in these pathways are reported below.

**Table 1.**
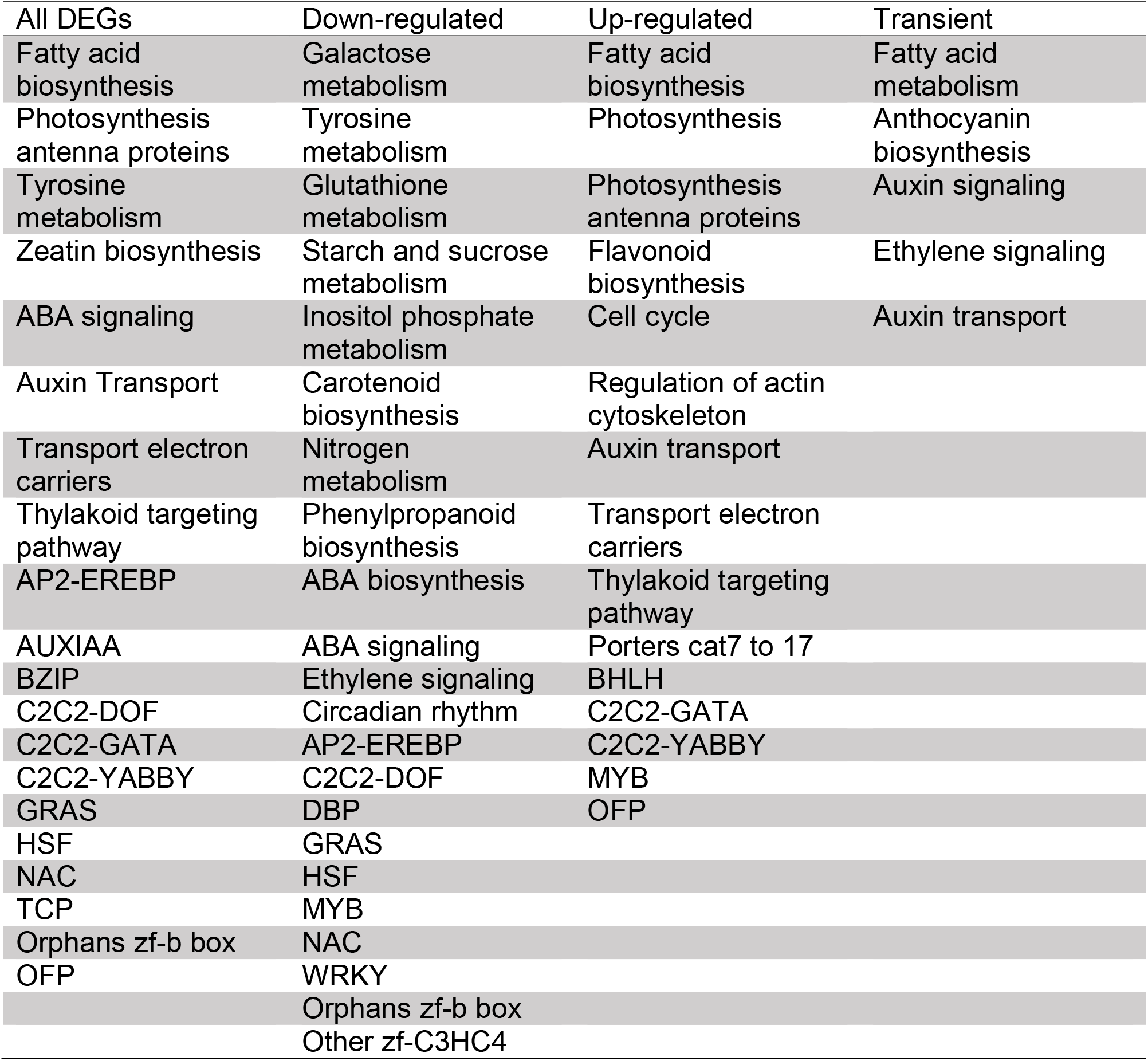
Enriched VitisNet pathways shared among *V. amurensis, V. riparia*, and *V. vinifera* ‘Cabernet Sauvignon’ and ‘Riesling’ during deacclimation and budbreak. Pathway enrichment analysis was conducted in VitisPathways using all differentially expressed genes (DEGs), as well as DEGs in the up-regulated, down-regulated, and transient expression groups.

Within the ABA biosynthesis pathway, zeaxanthin epoxidase (ZEP), 9-cis-epoxycarotenoid dioxygenase (NCED), and (ABA8’OH) were down-regulated while ABA glucosidase (ABA BG), was strongly up-regulated (Fig. 3). ABA signaling genes, such as *ABA-RESPONSIVE ELEMENT BINDING PROTEIN 2* (*AREB2*) and *KEEP ON GOING* (*KEG*), were down-regulated. Timing of *AREB2* downregulation was synchronized between genotypes, while the *KEG* was reduced faster in wild species than in *V. vinifera*. Both *ABA INSENSITIVE 1* (*ABI1*) and *ABA REPRESSOR 1* (*ABR1*) appear to have transient expressions. *MEDIATOR OF ABA-REGULATED DORMANCY 1* (*MARD1*) and Phospholipase D (PLD) genes are both generally up-regulated.

**Fig. 3.**
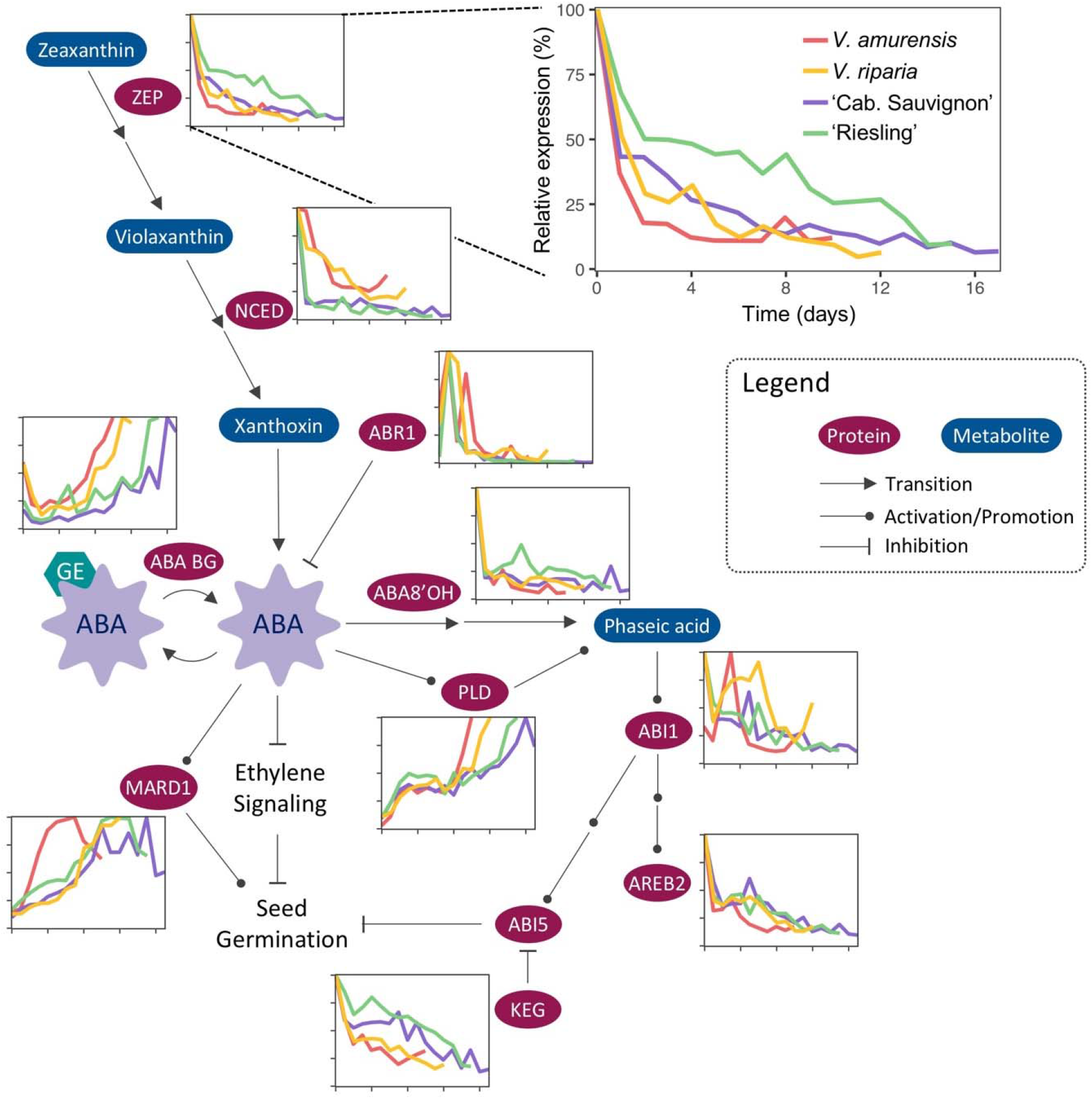
Reduced VitisNet ABA biosynthesis and signaling pathway. Mini-graphs indicate relative level of expression of any given protein for *V. amurensis* (red), *V. riparia* (yellow), and *V. vinifera* ‘Cabernet Sauvignon’ (purple) and ‘Riesling’ (green) during deacclimation and budbreak. Multiple DEGs encoding for the same protein had their expression aggregated. Multiple arrows indicate transition or signaling steps omitted. Detail shows expanded graph for axis information.

Ethylene synthesis was transiently up-regulated within a few days of deacclimation. 1-aminocyclopropane-1-carboxylate oxidase (ACC oxidase) and ACC synthase had peak expression after placement in warm conditions, or a temporary up-regulation (Fig. 4). Ethylene synthesis repressor gene *E8*, however, was up-regulated in all genotypes. Ethylene signaling, which includes ERFs, WRKY, and HSF pathways were primarily down-regulated (Supplementary Fig. S3, S4).

**Fig. 4.**
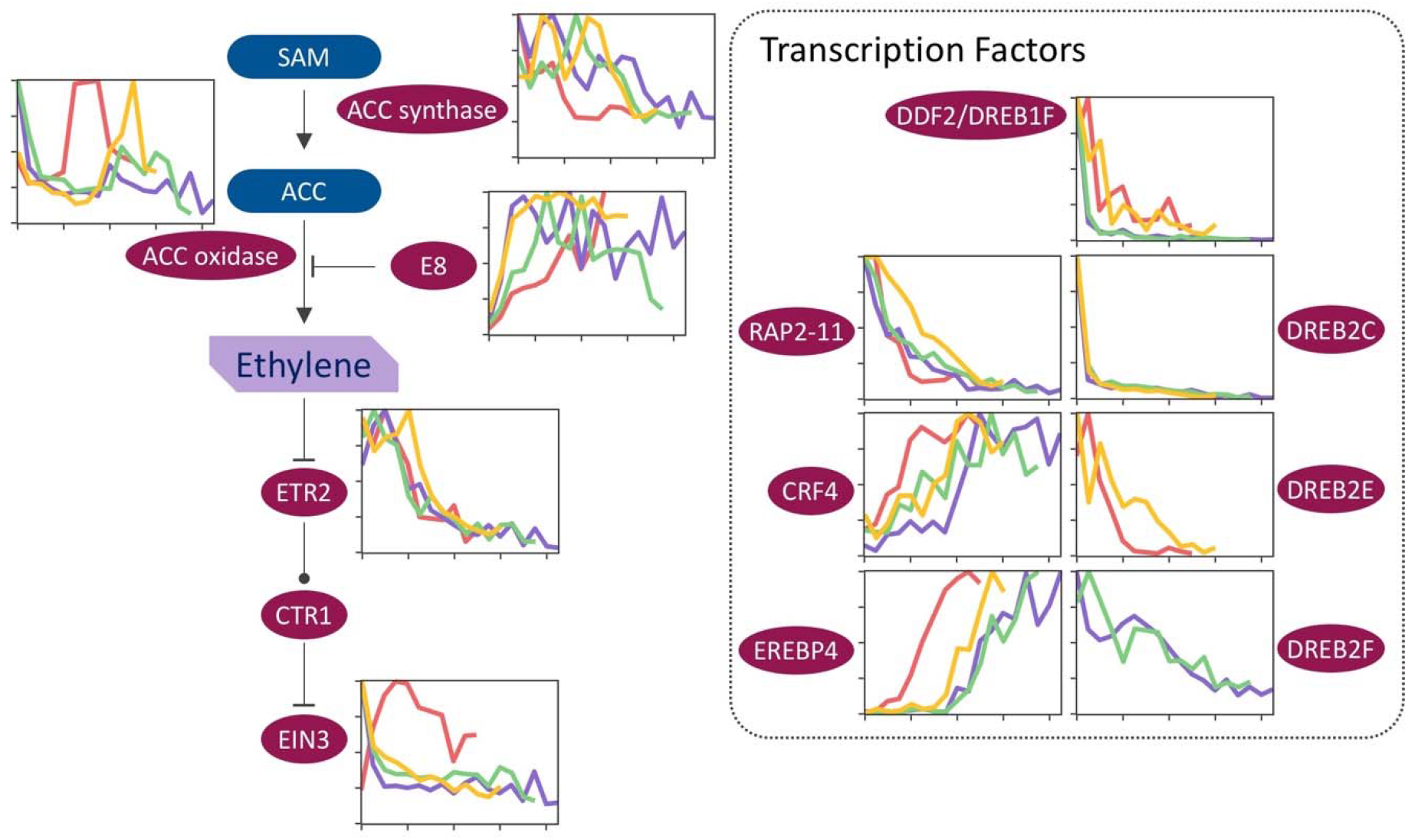
Reduced VitisNet ethylene biosynthesis and signaling pathway, and ethylene response factors (ERFs). Mini-graphs indicate relative level of expression of any given protein for *V. amurensis* (red), *V. riparia* (yellow), and *V. vinifera* ‘Cabernet Sauvignon’ (purple) and ‘Riesling’ (green) during deacclimation and budbreak. Multiple DEGs encoding for the same protein had their expression aggregated. Arrows indicate transition; circular arrows indicate positive regulation; and “T” indicates negative regulation. For details on graph axes and symbols see Fig. **3**.

Key components of jasmonate and gibberellin (GA) synthesis pathways were observed (Supplementary Fig. S5). Jasmonate synthesis was down-regulated, with precursors being led to medium chain fatty acids via up-regulation of lipoxygenase. MEJAE, which converts methyl-jasmonate to jasmonic acid, was down-regulated, whereas Jasmonate-O-methyltransferase, which catalyzes the opposite reaction, was up-regulated. For GAs, GA-3 β-dioxygenase is up-regulated in all genotypes, while GA-2 β-dioxygenase is down-regulated, indicating a trend towards synthesis of bioactive gibberellins. For Auxins, indole-3-acetic acid-amino acid hydrolases (IAA-AA-hydrolases), which reactivates IAA, was up-regulated in all genotypes, reaching maximum expression earlier in *V. amurensis* than other genotypes (Supplementary Fig. S6). *IAA6, IAA19, AUXIN RESPONSE FACTOR 5* (*ARF5*) and *AUXIN-RESISTANT 1* (*AUX1*) were up-regulated, with relative expression increasing faster in *V. amurensis* than other genotypes. Auxin responsive *PIN-FORMED* (*PIN*) *1* was down-regulated over time for all genotypes, while *PIN3* was up-regulated.

The degradation of cytokinin through cytokinin dehydrogenase (CKX) was up-regulated for all genotypes over time (Fig. 5). In addition, *PURINE PERMEASE 1*, a cytokinin transporter, was down-regulated. *AUTHENTIC RESPONSE REGULATOR* (*ARR*) *11* was down-regulated, and *ARR17* was up-regulated, further indicating a decrease in cytokinin content. Cyclin D, which is repressed by cytokinin responses and is the first cyclin involved in the cell cycle, was up-regulated earlier compared to cyclins A and B for all genotypes (Fig. 5). In fact, cyclins A and B have near 0 expression levels before cyclin D is up-regulated. Genes related to cell expansion and cell division (cyclins and cytoskeleton proteins) were up-regulated and staggered in accordance to species: *V. amurensis*, followed by *V. riparia*, and *V. vinifera* (Fig. 5).

**Fig. 5.**
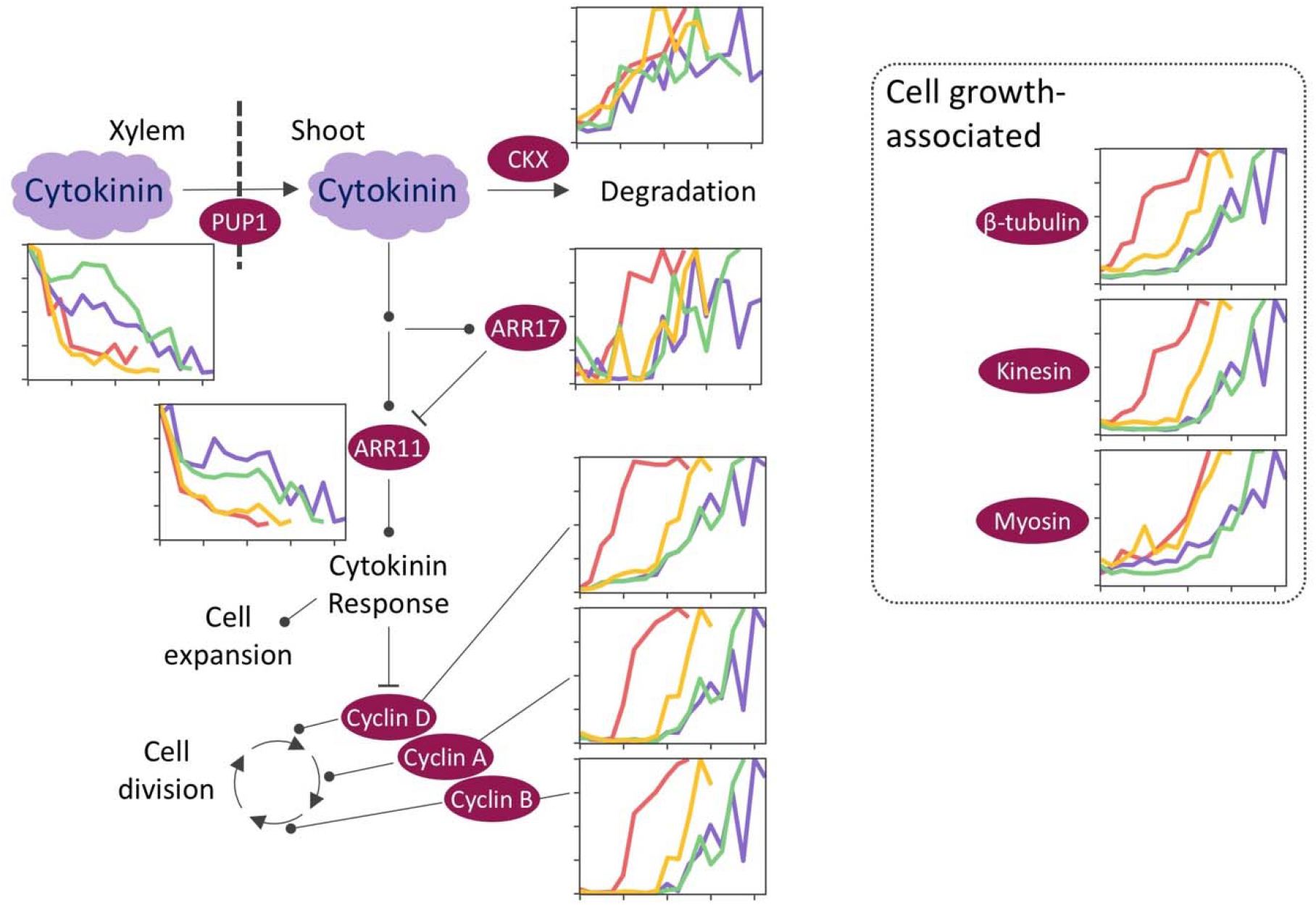
Reduced VitisNet cytokinin biosynthesis and signaling pathway, and growth associated genes. Mini-graphs indicate relative level of expression of any given protein for *V. amurensis* (red), *V. riparia* (yellow), and *V. vinifera* ‘Cabernet Sauvignon’ (purple) and ‘Riesling’ (green) during deacclimation and budbreak. Multiple DEGs encoding for the same protein had their expression aggregated. Arrows indicate transition; circular arrows indicate positive regulation; and “T” indicates negative regulation. Multiple arrows indicate transition or signaling steps omitted. For details on graph axes and symbols see Fig. **3**.

Circadian rhythm pathway genes were generally down-regulated in all genotypes (Supplementary Fig. S7). *CIRCADIAN 1* (*CIR1*), *EARLY FLOWERING 3* (*ELF3*), and *ARABIDOPSIS PSEUDO-RESPONSE REGULATOR 5* (*APRR5*) had a strong down-regulation in the first few days, whereas *CIRCADIAN CLOCK ASSOCIATED 1* (*CCA1*) and Catalase *CAT3* reduced at a slower rate. In the phenylpropanoid biosynthesis pathway, phenylalanine ammonia lyase was initially down-regulated, followed by up-regulation that occurs earlier in *V. amurensis*, followed by *V. riparia*, and the *V. vinifera* genotypes (Supplementary Fig. S8). Trihydroxystilbene synthase (stilbene synthase) was quickly down-regulated in all genotypes, but faster in the wild species than *V. vinifera*. Of the 44 predicted genes for stilbene synthase, between 29 and 35 were DEGs for down-regulation in all genotypes.

Sugar metabolism genes encoding Starch and Glycogen [starch] synthases, and beta amylase were down-regulated (Fig. 6). However, Glycogen [starch] synthase and beta amylase are synchronized in all genotypes, whereas starch synthase reduces its expression earlier in wild genotypes. Sucrose synthase is initially down-regulated in all genotypes, while a return to high levels of expression is seen in *V. riparia* and *V. vinifera*, but not in *V. amurensis*. Fructokinase and hexokinase are generally up-regulated in all genotypes, leading sugars into glycolysis. Similarly, genes in fatty acid biosynthesis and metabolism pathways were generally up-regulated over time (Supplementary Fig. S9), including those involved in β-oxidation. *FATTY ACID DESATURASE 5* (*FAD5*), however, was strongly down-regulated for all genotypes.

**Fig. 6.**
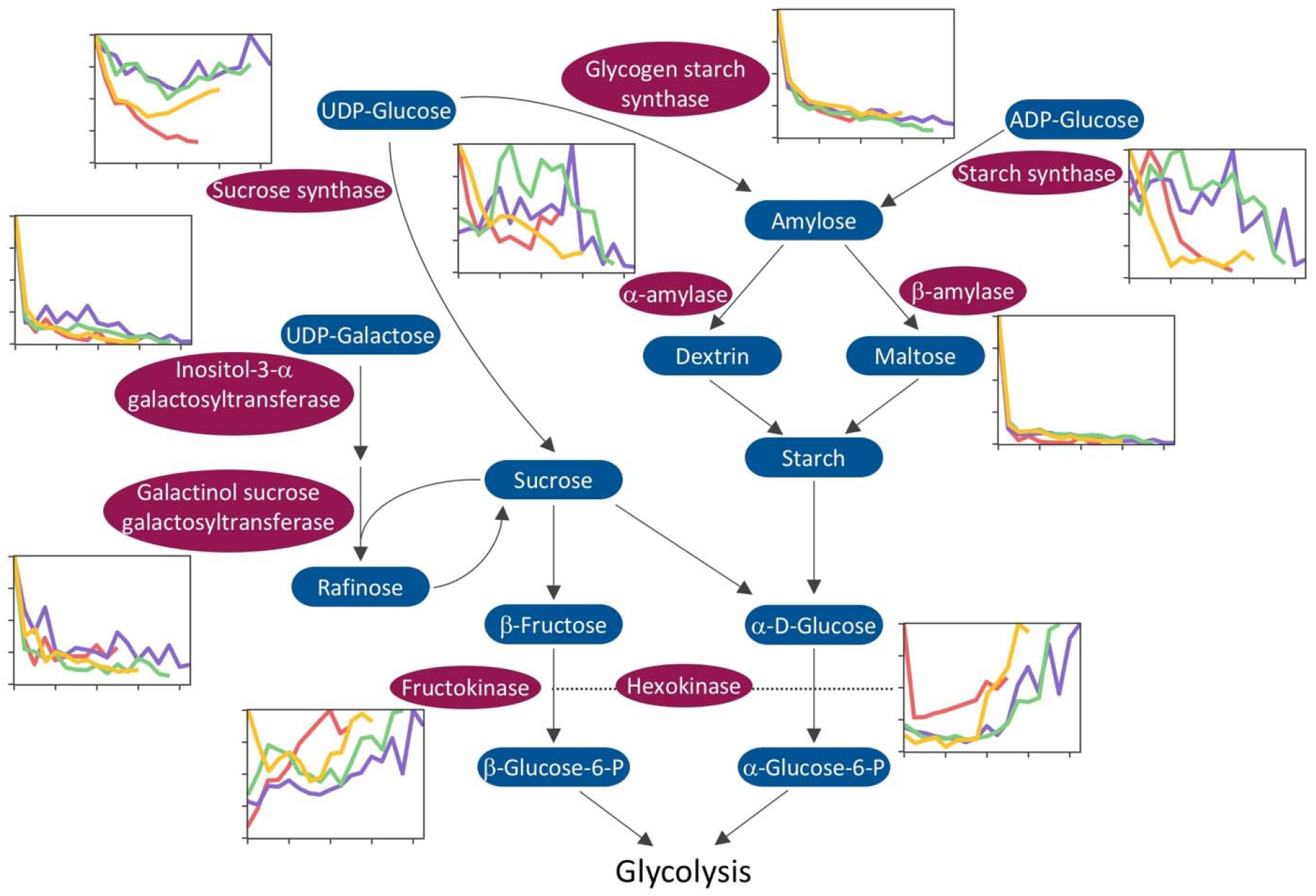
Reduced VitisNet starch, sucrose, and galactose metabolism pathways. Mini-graphs indicate relative level of expression of any given protein for *V. amurensis* (red), *V. riparia* (yellow), and *V. vinifera* ‘Cabernet Sauvignon’ (purple) and ‘Riesling’ (green) during deacclimation and budbreak. Multiple DEGs encoding for the same protein had their expression aggregated. Arrows indicate transition. Multiple arrows indicate transition steps omitted. For details on graph axes and symbols see Fig. **3**.

In photosynthesis related pathways, genes encoding light harvesting complex I (LHCI) and II (LHCII), and photosystem I (PSI) subunits also follow the trend of growth-related genes, with earlier up-regulation in wild species (Supplementary Fig. S10). *RESPIRATORY BURST OXIDASE HOMOLOG PROTEIN B* (*RBOHB;* all genotypes) and *RBOHE* (all but *V. riparia*) had transient up-regulation in the earlier days. *RBOHF* was also up-regulated in the first few days, returning to ~50% relative expression. Peak expression of *RBOHF* occurs a few days prior to budbreak in *V. amurensis, V. riparia*, and *V. vinifera* ‘Cabernet Sauvignon’. *RBOHD* appeared to have an increasing expression trend.

Vacuole localized aquaporins *TONOPLAST INTRINSIC PROTEIN* (*TIP*) *3;1* and *TIP3;2* were down-regulated, while *TIP1;3* and *TIP4;1* were up-regulated (Fig. 7). *CYCLIC NUCLEOTIDE-GATED ION CHANNEL PROTEIN 15* (*CNGC15*) was down-regulated in all genotypes.

**Fig. 7.**
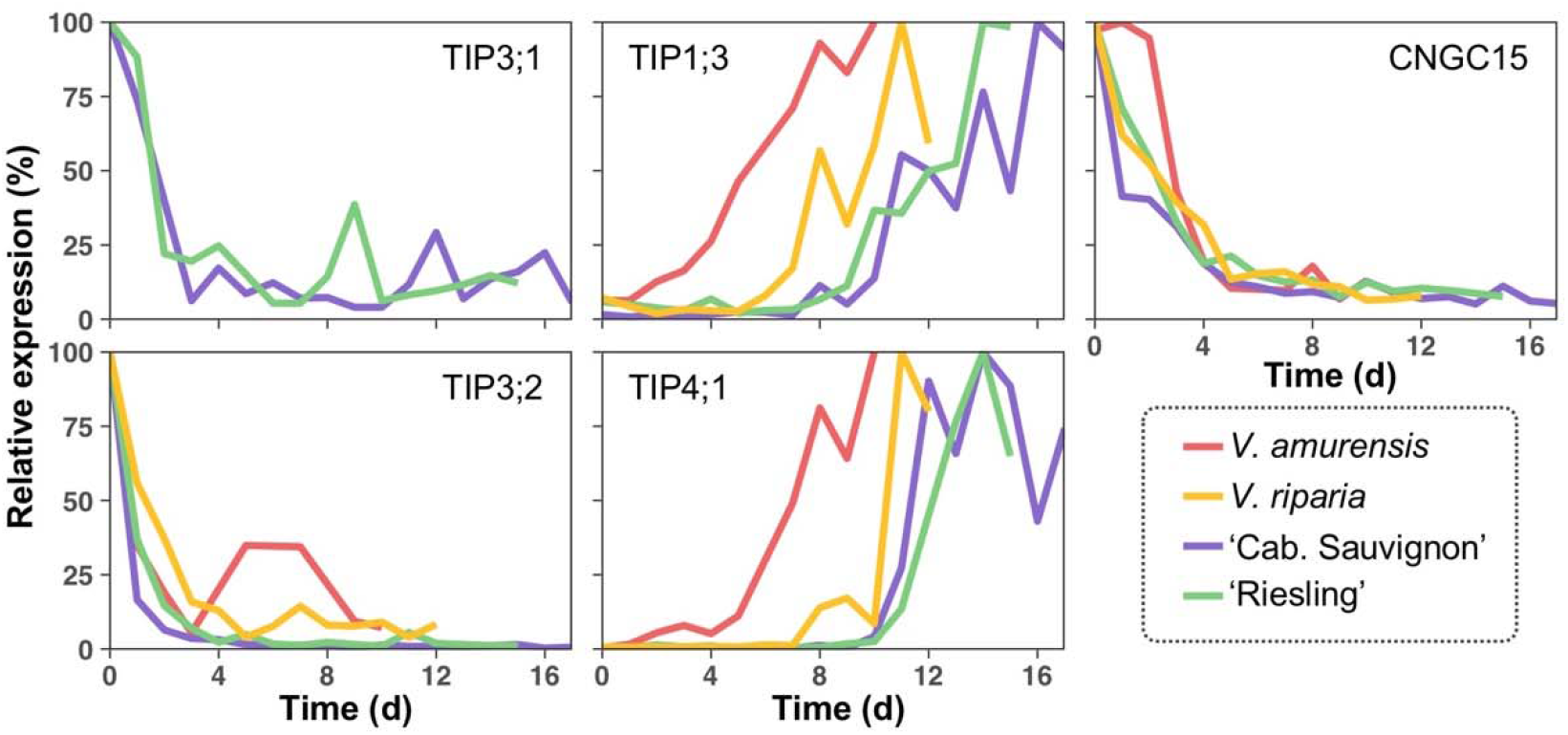
Relative expression of channel protein genes during deacclimation and budbreak in *V. amurensis* (red), *V. riparia* (yellow), and *V. vinifera* ‘Cabernet Sauvignon’ (purple) and ‘Riesling’ (green).

### ABA effects on cold hardiness

ABA treatments had no effect on budbreak or deacclimation until 13 days post application (Fig. 8A, B). Starting on day 16, ABA treatments prevented further loss of cold hardiness, while deacclimation continued in the water control (Fig. 8B). Both ABA concentrations maintained significantly more cold hardiness than the control, with no difference amongst themselves. After ABA treatment was removed, previously treated buds resumed deacclimation. Regarding phenology, control buds were significantly more developed starting on day 44 in pairwise comparisons with ABA treated buds, although buds in the control treatment average E-L stage > 1 starting on day 30 (Fig. 8A). Control buds reached budbreak (E-L stage 3) at 49.0±1.5 days after the beginning of the experiment as estimated by a logistic regression. For 1mM ABA, budbreak occurred at 73.2±3.3 days, while buds in 5mM did not resume growth before the experiment ended at 84 days.

**Fig. 8.**
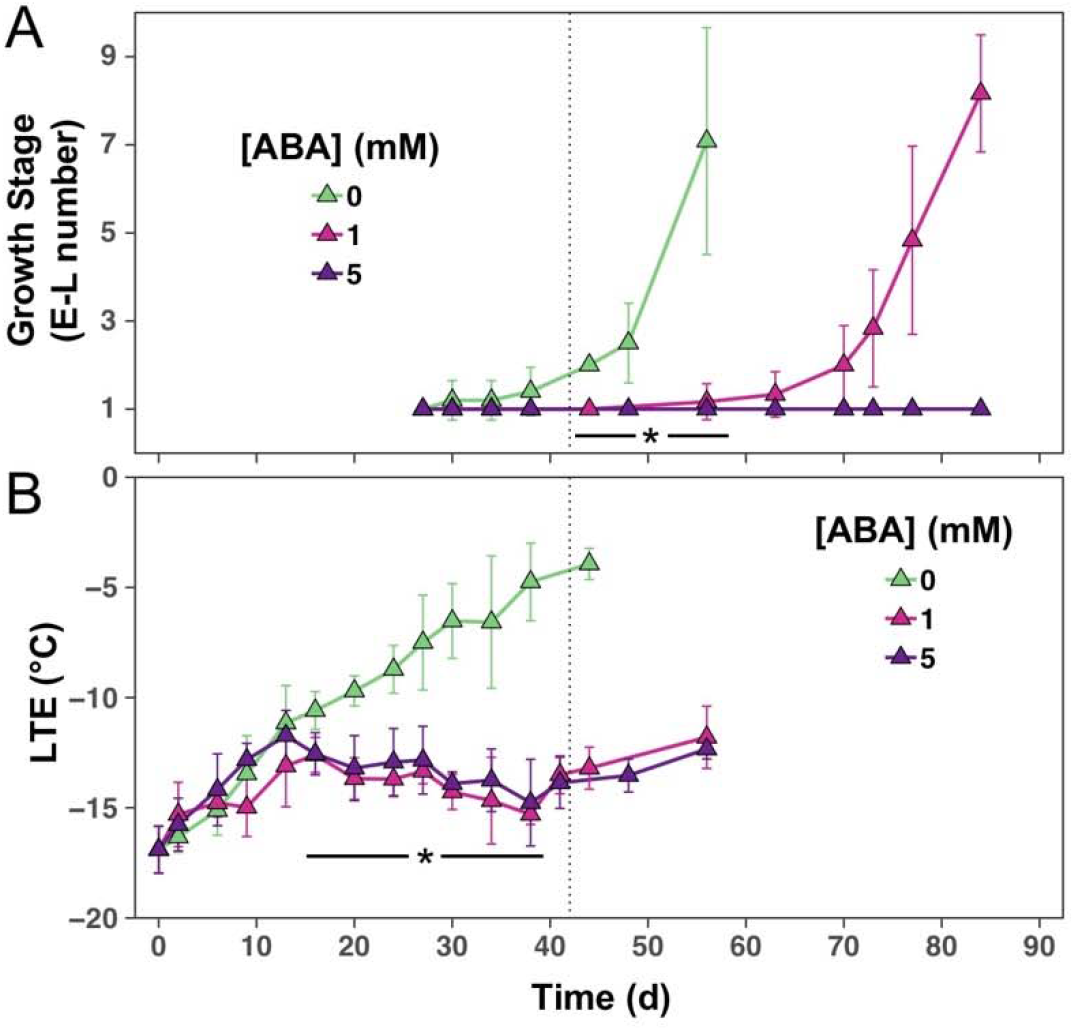
Cold hardiness and budbreak of *V. vinifera* ‘Riesling’ buds treated with abscisic acid (ABA). (a) Stage of development (E-L number; Coombe and Iland, 2005) and (b) cold hardiness of *V. vinifera* ‘Riesling’ buds in 0, 1, and 5 mM abscisic acid solution at an average temperature of 7 °C 0/24h light/dark photoperiod. After 42 days (vertical dotted line) samples were moved to 22 °C 16/8h light/dark photoperiod. Error bars represent standard deviations of the mean.

## Discussion

Deacclimation and growth are processes that occur in tandem in grapevine, although most or all of the cold hardiness is lost before growth is visible. As these processes occur incrementally over many days, a novel approach for time-series data analysis to find DEGs was used. To our knowledge, this is the first study exploring long-term RNA-Seq data for the exploration of genes and pathways important for deacclimation and budbreak. We used species with contrasting phenotypes to generate hypotheses and explore the genes that are likely related to the cold hardy/dormant phenotype, those related to growth and budbreak, and those that may be shared between both.

The approach developed for data analysis in this study was important to detect many trends in the long-term RNA-Seq data for differential expression. Here we treated time as a continuous variable, modeling counts with a fourth degree polynomial to allow transient expression of genes to be properly predicted, and using the predicted counts for filtering. Typically, studies will either use pairwise comparisons between adjacent timepoints (e.g., Meitha *et al*., 2018) or between each timepoint and a 0h control (e.g., Fu *et al*., 2018). The first approach may result in a low number of DEGs (Días-Riquelme *et al*., 2012), while the second only accounts for changes in gene expression that are different from a “basal” condition. Differences in time between sample points [e.g., 3h-72h (Meitha *et al*., 2018); 3h-120h (Fu *et al*., 2018)] also argues that time may be more appropriate as a continuous variable.

While the approach in Meitha *et al*. (2018) would more likely detect genes with transient expression or genes with extreme changes in expression, and the approach of Fu *et al*. (2018) might miss transiently expressed genes with maximum and minimum expression occurring after the first time point, our approach makes no assumption as to the timing of significant expression change.

Sampling within a 24h period can result in the detection of “false” DEGs due to circadian oscillations in genes (Grundy *et al*., 2015). In our case, samples were always taken within a 3h window. Irregular sampling could benefit from using time as a continuous variable and sine/cosine functions that may elucidate some of the variation in gene expression due to the circadian rhythm (Cronn *et al*., 2017). Using continuous variables for time for RNA-Seq studies with more than 3 data points might be the most appropriate method, and even time-series data with lower replicate numbers [e.g., 2 biological replicates (Sozzani *et al*., 2010)] could benefit from this type of analysis for more rigor than pairwise comparisons considering how degrees of freedom are distributed for continuous variables.

Our objective was to observe asynchronous behavior in gene expression relative to asynchronicity of the deacclimation pattern, therefore we analyzed the data in response to time within each genotype. It is clear that correcting gene expression data by inherent phenological development rates may be necessary for other experiments (Fig 2A, B, C). The results presented here demonstrate that not only does phenology in grapevine correlate with deacclimation kinetics (Kovaleski *et al*., 2018), so do the patterns of gene expression that occur during this physiological transition. As wild grapevines are used in cold hardy hybrid cultivar breeding, it is important to recognize that these cold hardy genotypes also contribute faster deacclimation and budbreak phenology (Fig 1A; Londo and Kovaleski, 2017; Kovaleski *et al*., 2018) controlled by the accelerated expression of the deacclimation cascade (Supplementary Fig. S2).

Temperature sensing must be the first step directing the physiology of the bud towards growth. Changing the environment from field to growth chamber (−7 °C to 22 °C) between day 0 and day 1 resulted in rapid temperature sensing-related transcriptional changes. Three lines of evidence for this rapid temperature sensing were observed. Membrane rigidity has long been suggested as the most upstream temperature sensor for thermal signaling (Horváth *et al*., 1998, 2012; Saidi *et al*., 2010; Bahuguna and Jagadish, 2015). Our results agree with this hypothesis as fast down-regulation of *FAD5* and general up-regulation of fatty acid biosynthesis genes (Supplementary Fig. S9) likely lead to increased content of saturated fatty acids and decreased membrane fluidity (Wallis and Browse, 2002; Falcone *et al*., 2004; Filek *et al*., 2017). CNGCs are also important in thermosensing as channel proteins for Ca^2+^ signaling, and have been implicated in dormancy release of buds (Pang *et al*., 2007; Bahuguna and Jagadish, 2015). Although plasma membrane localized CNGCs have clear roles in cold sensing (Bahuguna and Jagadish, 2015), here we observed down-regulation of *CNGC15* [nucleus localized (DeFalco *et al*., 2016)] was detected in all genotypes. This suggests that in grapevine buds, Ca^2+^ stored in the nuclear envelope lumen or endoplasmic reticulum may contribute to signaling during the dormant period. Finally, PLD expression increased in response to dormancy release. PLD has been observed to be up-regulated during cold and freezing stress (Ruelland *et al*., 2002) as well as many other stress conditions (Meijer and Munnik, 2003), and is indicated as a signaling protein in thermal sensing (Bahuguna and Jagadish, 2015). PLD activity is also positively influenced by Ca^2+^ influx during cold stress (Meijer and Munnik, 2003; Wang and Nick, 2017). Overexpression of PLD results in higher freezing tolerance (Li *et al*., 2004), and more rapid and sensitive response to ABA and stress (Sang *et al*., 2001; Meijer and Munnik, 2003). Considering our treatment can be seen as a release from cold stress, but PLD expression increased, we speculate that effects of PLD in cold stress-relief signaling are likely related to changes in plasma membrane composition through selective hydrolysis of membrane phospholipids (Wang, 1999).

Following sensing of warm temperatures, we expected a decrease in cold-related transcription factors would occur. Interestingly, none of the *CBF/DREB1* genes were differentially expressed, and within *DREB1s* only *DREB1F/DDF2* was differentially expressed. It is possible that CBFs are involved in cold hardiness gain and in response to cold stress (Liu *et al*., 2017a; Ban *et al*., 2017; Rubio *et al*., 2018), but are not maintained in a high level of transcription throughout the winter. Alternatively, DREB2 homologs were differentially expressed and down-regulated. This result disagrees with Liu *et al*. (1998) and Nguyen *et al*. (2017), who suggested *DREB1* and *DREB2* are two independent *DREB* families that lead to signal transduction pathways for low temperature and dehydration conditions, respectively. Consequently, when buds were moved into forcing conditions, down-regulation of *CBF/DREB1s* transcripts was not detected. Instead, lack of vascular continuity in early stages of budbreak (Xie *et al*., 2018) may result in some level of bud drought stress, contributing to the DREB family patterns observed.

Pathways and genes associated with stress were downregulated, such as ABA, ethylene, ROS burst and detoxification, and circadian clock genes. ABA biosynthesis is a requirement for cold acclimation and thermotolerance (Gilmour and Thomashow, 1991; Larkindale *et al*., 2005; Penfield, 2008), and ABA is a clear antagonist of deacclimation and growth. Both the ABA biosynthesis and the signaling pathways were down-regulated in this study. The pivotal role of the ABA synthesis reduction for the loss of hardiness is clear (Fig. 3). Both ZEP and especially NCED, the rate-limiting enzyme in ABA biosynthesis (Grundy et al., 2015), were quickly down-regulated in all genotypes, indicating carotenoids are likely no longer contributing to synthesis of new ABA during deacclimation. ABR1, an AP2 TF repressor of ABA response (Pandey *et al*., 2005), was quickly up-regulated from day 0 to the highest relative level of expression in all genotypes at day 1, indicating the move to warm temperatures quickly signals for reduction of ABA synthesis as well as repression of ABA responses from endogenous pools. Supporting the role of ABA in reinforcing dormancy, high exogenous ABA concentrations halted deacclimation (Fig. 8), although there was no net gain in hardiness as in Rubio *et al*. (2018). ABA BG expression was upregulated while the rest of the synthesis and signaling pathway was downregulated, suggesting that reactivation of inactive vacuolar ABA may play a role in preventing dormancy release during endodormancy. Given there was no lag in deacclimation response it is unlikely that much vacuolar storage of inactive ABA remained when our study was conducted. Future studies should separately examine the fates and concentrations of newly synthesized ABA and reactivated ABA during deacclimation, and determine if these separate pools could be leveraged as a biomarker for dormancy status. Another important role of ABA during dormancy appears to be in controlling water status in the bud, which is critical for low temperature survival (George and Burke, 1977). TIP3;1 and TIP3;2 – ABA regulated aquaporins (Mao and Sun, 2015) – were down regulated as opposed to the general up-regulation of TIP aquaporins (Fig. 7). While other aquaporins have been implicated in chill stress responses (Bilska-Kos *et al*., 2016), regulation of TIP3s in our study demonstrates that ABA may be affecting water movement in the bud during the dormant season, maintaining cold hardiness.

Steps leading to ethylene biosynthesis appear transiently up-regulated in the initial days of warm period in this study. This may be due to quick loss of ABA’s antagonistic effect on the synthesis of ethylene. Ethylene, along with H_2_O_2_, lead to activation of antioxidative stress genes in grapevine (Vergara *et al*., 2012). H_2_O_2_ is involved in the regulation of cold-responsive genes expression during acclimation (Fedurayev *et al*., 2018), during forced budbreak with hydrogen cyanamide (Or *et al*. 1999; Sudawan *et al*. 2016), and temporary increases in ROSs have been described to precede budbreak (Ophir *et al*., 2009; Pérez and Lira, 2005; Khalil-Ur-Rehman *et al*., 2017; Meitha *et al*., 2018), demonstrating contrasting roles. RBOHF is involved in ethylene and ABA signaling through H_2_O_2_ production (Kwat *et al*., 2003; Desikan *et al*., 2006), and the expression pattern in our study appears to be a marker for budbreak. While gene expression in relation to hypoxia and ethylene have been previously described during budbreak (Ophir *et al*., 2009; Meitha *et al*., 2018), this study identified a potentially novel role of stilbene synthase. Stilbene synthase is responsible for resveratrol production in the phenylpropanoid pathway, which is an ROS scavenger. The down-regulation of the majority of the 44 genes encoding stilbene synthase in all 4 genotypes after collection from the field suggests resveratrol production is an important part of cold hardiness or dormancy maintenance. Stilbene synthase has previously been described as up-regulated in grapevine leaves during cold or freeze stress (Londo *et al*., 2018), but is also involved in other types of stress during the growing season (Carvalho *et al*., 2015). Similarly, studies examining cold tolerant *Passiflora edulis* (Liu *et al*., 2017a) and ecodormant *Pyrus pyrifolia* (Bai *et al*., 2013) support the hypothesis of phenylpropanoids as cold hardiness and dormancy metabolites.

Regulation of circadian clock genes is important for abiotic stress tolerance (Grundy *et al*., 2015), and alternative splicing has been indicated as a mechanism of signaling in cold response (Calixto *et al*., 2016). Most genes in the circadian rhythm pathway were down-regulated over time. Both *CCA1* and its promoter *CIR1* are down-regulated in all genotypes. *CCA1* is a transcription factor that regulates low temperature responses through control of *CBF* gene expression (Grundy *et al*., 2015), and also represses *TOC1* expression. Higher expression of *CCA1* in the field may be disrupting of the circadian clock (Ramos *et al*., 2005), but may also be required during dormancy in order to maintain the circadian rhythm without strong effects of day length or light.

Growth and cell expansion pathways were upregulated rapidly and sequentially after day 1. The trend towards synthesis of bioactive GA suggested by down-regulation of GA-2 β-dioxygenase and up-regulation of GA-3 β-dioxygenase (Supplementary Fig. S5) was also reported by Bai *et al*. (2013) comparing eco- and endodormant buds. Those authors also observed an up-regulation of Jasmonate-O-methyltransferase as seen here (Supplementary Fig. S5). This interplay between MEJAE and Jasmonate-O-methyltransferase suggest that methyl-jasmonate is the preferred form of jasmonate during budbreak.

Genes in carbohydrate metabolism likely have significant roles during the dormancy period in grapevine buds (Khalil-Ur-Rehman *et al*., 2017). While Starch and Sucrose Metabolism pathway is enriched with down-regulated genes (Table **1**), and enzymes in the degradation of starch are generally down-regulated, fructokinase and hexokinase are both up-regulated (Fig. 6). This indicates that pools of sugar are being used in glycolysis to generate energy, but also that sucrose used in sugar metabolism is being sourced outside the bud. The up-regulation of Enoyl CoA hydratase and Acetyl CoA c-Acyltransferase in the fatty acid metabolism (Supplementary Fig. S9) indicates that energy is also being produced from free fatty acids such as those resulting from the degradation of phospholipids by PLD (Wang, 1999).

The same temporal staggering observed for the budbreak phenotype was expected to be reflected in gene expression patterns for genes related to growth. This correlation is clearly observed in the up-regulation of cyclins and cytoskeleton proteins in all genotypes (Fig. 5), following observed timing of budbreak (Fig. 1B). Within genotypes, earlier expression of cyclin D indicates that the G_1_ and S phases of the cell cycle (Mironov *et al*., 1999) occur concomitantly to deacclimation. However, the initially very low expression of both cyclin A and B indicates that G_2_ and M phases only occur after cold hardiness is lost, suggesting deacclimation and budbreak are cell expansion-but not cell division-dependent, and expression of these genes may be used as markers for imminent budbreak.

What determines the phenotype difference between fast and slow deacclimators? It appears that both sensing of the stimulus for dormancy release (change in environment) and down-regulation of cold-hardiness- and dormancy-maintaining genes occurred simultaneously between all genotypes. This is observed in the synchronous down-regulation of *CNGC15, FAD5*, ABA synthesis, and stilbene synthase, among others. Differences between the four genotypes in both phenotypes observed (deacclimation and budbreak) must therefore arise from how quickly genotypes are able to re-establish growth. This is exemplified in the staggering of expression seen in cyclins, cytoskeleton- and auxin-related genes. Many players have been implicated in the control of growth, but here we indicate those that may also play a role in deacclimation, or promote growth through promoting deacclimation. We thus propose a hypothetical model for events occurring during release of dormancy in grapevine buds (Fig. 9). Upon sensing of growth permissive temperatures, likely by membrane fluidity, ABA synthesis is down-regulated. With the release of inhibition by ABA, a temporary increase in ethylene synthesis and signaling leads to a transient burst of ROS. An influx of water and cell expansion slowly increase water potentials, decreasing cold hardiness. At the same time, energy is generated from fatty acids and cane-stored carbohydrates, and growth hormone pathways are up-regulated. Ultimately, with cell expansion the appearance of budbreak occurs, along with the start of cell division. As tissues emerges from buds, photosynthesis-related proteins are ready to start generate energy – although the sink to source switch in breaking buds occurs beyond the time analyzed here.

**Fig. 9.**
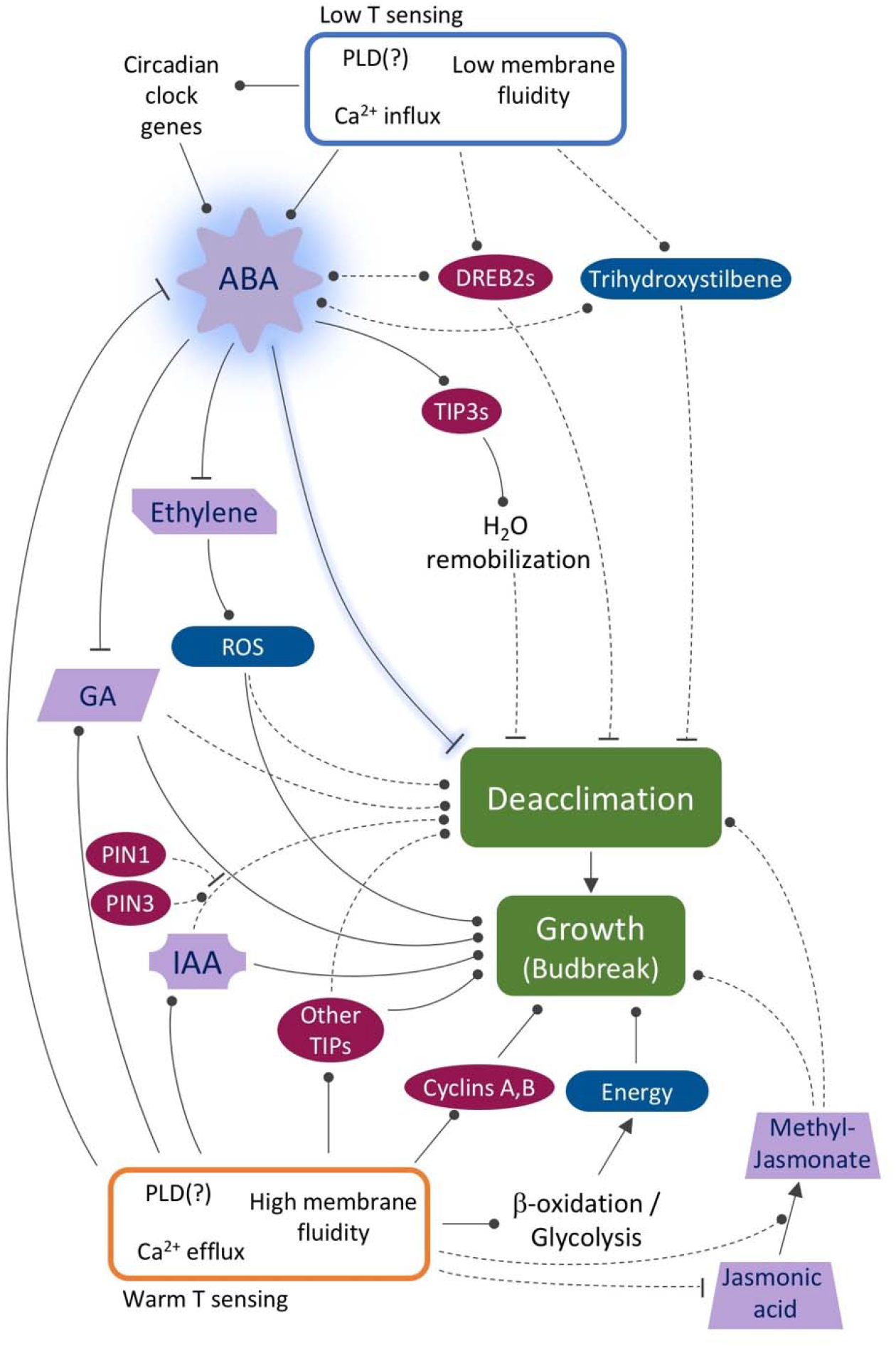
Schematic representation of proposed molecular control of deacclimation and growth based on gene expression data from this study. ABA appears as a master regulator of cold hardiness and dormancy maintenance, inhibiting commonly known routes that promote budbreak. Arrows indicate transition; circular arrows indicate positive regulation; and “T” indicates repression. Solid lines indicate validated interactions. Role of ABA in repression of deacclimation observed here highlighted in blue. Dashed lines are new interactions supported by gene expression data.

Understanding physiological transitions such as deacclimation and budbreak are essential for improving crop sustainability in the face of changing climate. We identify important genes and pathways related to maintaining cold hardiness, the loss of cold hardiness (deacclimation), the release of dormancy and budbreak by exploring phenotypic or temporal differences between *Vitis* spp. genotypes. It is likely that processes described here are conserved in other perennial woody species which show similar budbreak behavior in response to chilling [e.g., *Prunus persica* (Fan *et al*., 2010)]. In this study, we describe a new way to deal with variable length, time-series data such that a single list of DEGs is produced from all time-points. In our discussion we demonstrate that molecular processes involved in gain of cold hardiness are not just in reverse mode during deacclimation (e.g., PLD regulation, CBFs). ABA and its signaling were confirmed to be closely downstream from temperature sensing, and upstream from what defines cold hardiness with exogenous ABA. Therefore, ABA has the potential for mitigation of unseasonal or early deacclimation and budbreak in warmer winters. Finally, we discovered differences in deacclimation rate and initial cold hardiness contribute to variation in budbreak phenology between genotypes through differential gene regulation. Implications of this association may directly link to spring phenology markers and improved budbreak prediction in agricultural conditions as they experience climate variation.

## Supplementary Data

**Fig. S1**. Venn diagrams of shared genes.

**Fig. S2**. Heatmap of top 1000 genes.

**Fig. S3**. WRKY transcription factors.

**Fig. S4**. HSF transcription factors.

**Fig. S5**. Jasmonate and gibberellin biosynthesis pathways.

**Fig. S6**. Auxin biosynthesis pathway and auxin-related genes.

**Fig. S7**. Circadian rhythm pathway.

**Fig. S8**. Phenylpropanoid biosynthesis pathway.

**Fig. S9**. Fatty acids synthesis and metabolism.

**Fig. S10**. Photosynthesis-related genes.

## Acknowledgements

This work was partially supported by CAPES, Coordenação de Aperfeiçoamento de Pessoal de Nível Superior, Brazil, award number 12945/13-7, and by U.S. Department of Agriculture appropriated project 1910-21220-006-00D.

