## Supplementary Figures S1-S10 for "Tempo of gene regulation in wild and cultivated *Vitis* species shows coordination between cold deacclimation and budbreak"

**Fig. S1** Venn diagrams of shared differentially expressed genes (DEGs) between *Vitis amurensis* (red), *V. riparia* (yellow), and *V. vinifera* 'Cabernet Sauvignon' (purple) and 'Riesling' (green) during deacclimation and budbreak. Venn diagrams were produced using (A) all DEGs, (B) down-regulated, and (C) up-regulated genes.

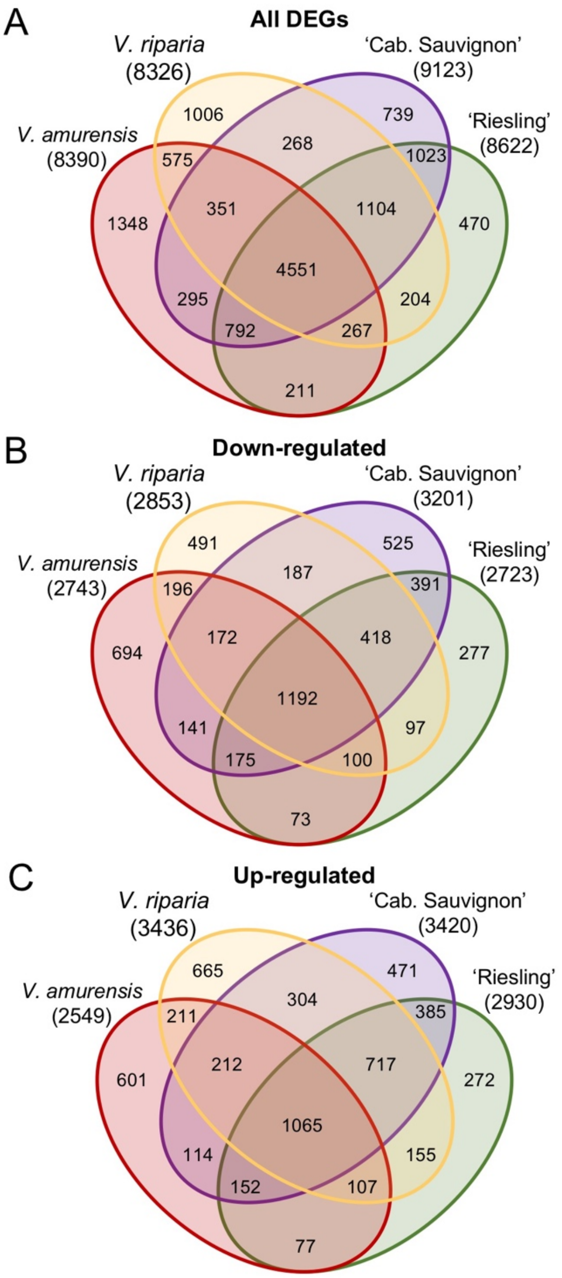

**Fig. S2** Heatmap of normalized counts (log2CPM) of top 1000 row variance genes for *Vitis amurensis* (red), *V. riparia* (yellow), and *V. vinifera* 'Cabernet Sauvignon' and 'Riesling' over time and thermal time (DDD) during deacclimation and budbreak.

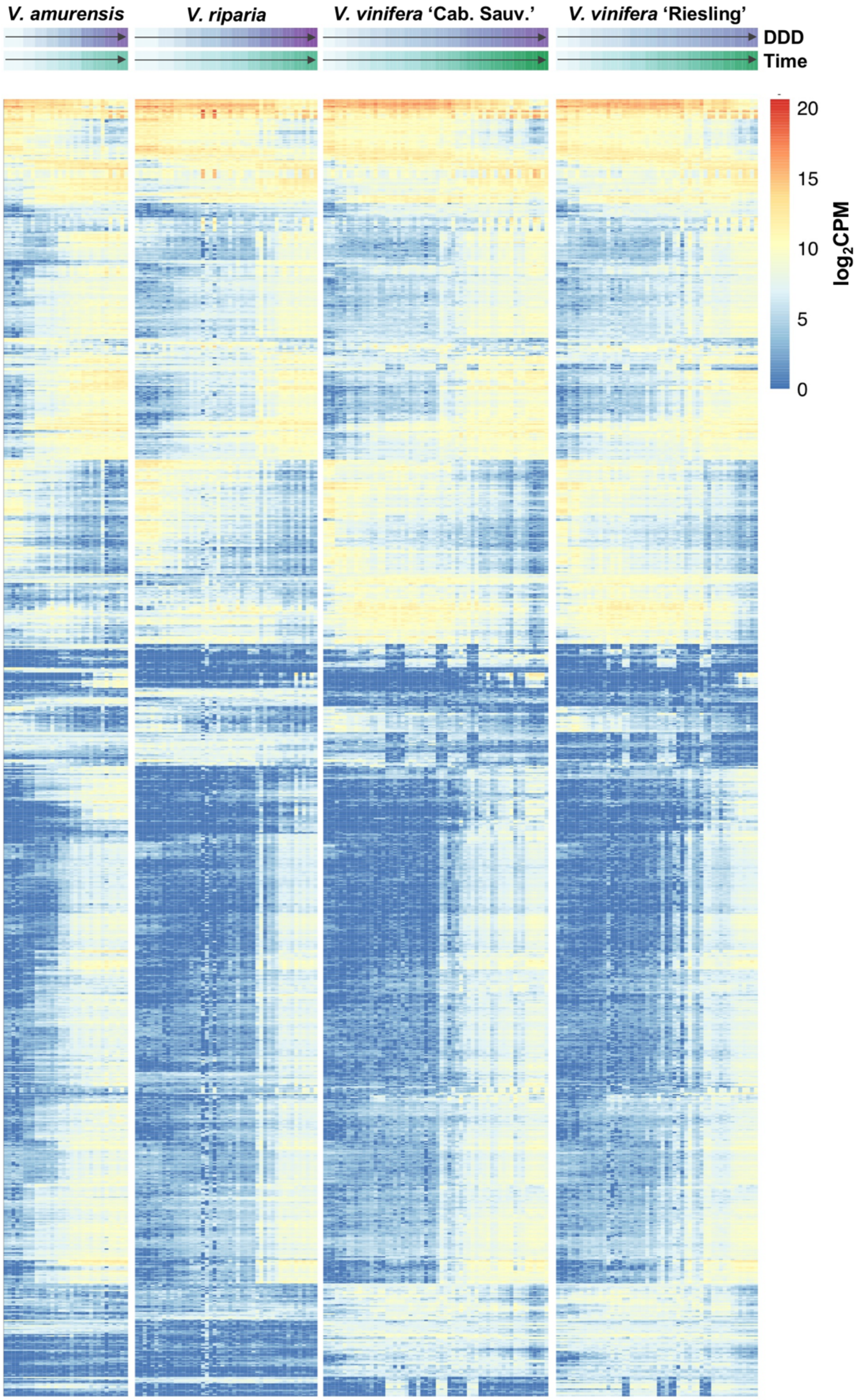

**Fig. S3** Relative expression of genes in the WRKY family of transcription factors during deacclimation and budbreak in *Vitis amurensis* (red), *V. riparia* (yellow), and *V. vinifera* 'Cabernet Sauvignon' (purple) and 'Riesling' (green).

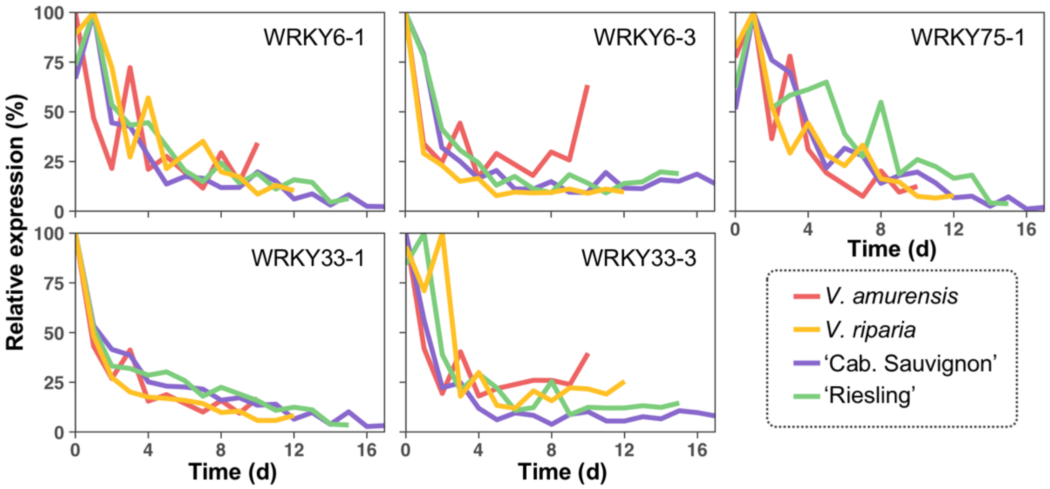

**Fig. S4** Relative expression of genes in the HSF family of transcription factors during deacclimation and budbreak in *Vitis amurensis* (red), *V. riparia* (yellow), and *V. vinifera* 'Cabernet Sauvignon' (purple) and 'Riesling' (green).

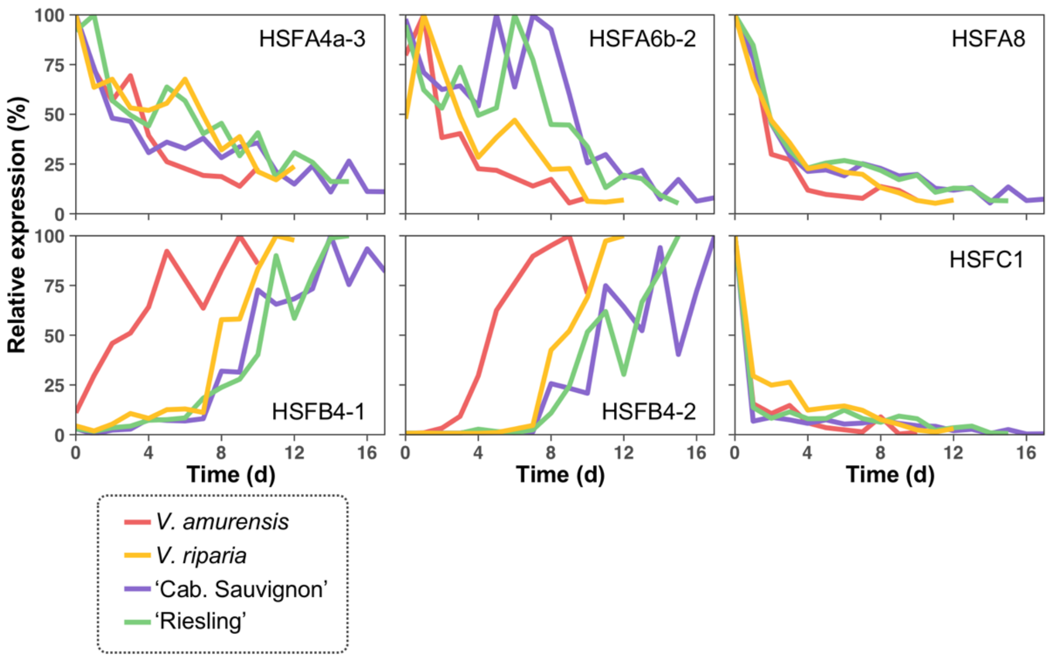

**Fig. S5** Reduced VitisNet Jasmonic Acid and Gibberellin biosynthesis pathways. Mini-graphs indicate relative level of expression of any given protein for *Vitis amurensis* (red), *V. riparia* (yellow), and *V. vinifera* 'Cabernet Sauvignon' (purple) and 'Riesling' (green) during deacclimation and budbreak. Multiple DEGs encoding for the same protein had their expression aggregated. Arrows indicate transition. Multiple arrows indicate transition steps omitted. For details on graph axes and symbols see Fig. **3**.

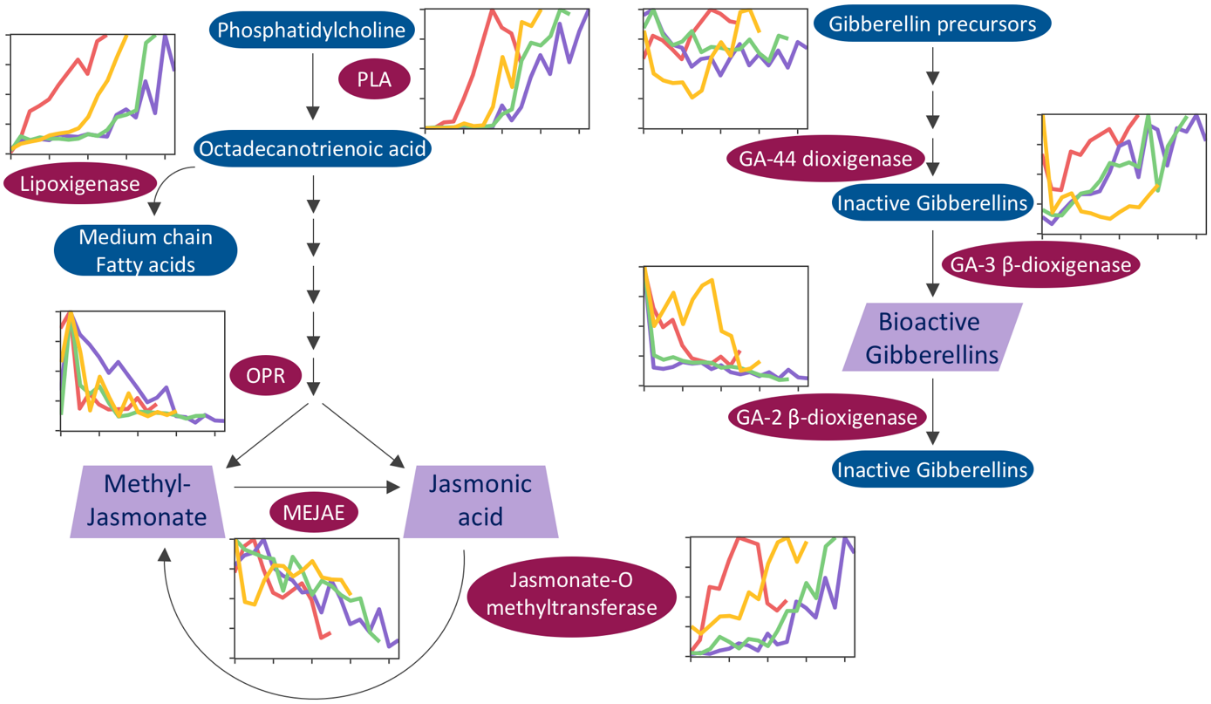

**Fig. S6** Reduced VitisNet auxin biosynthesis pathway and auxin-regulated transcription factors (TFs) and transport-related genes. Mini-graphs indicate relative level of expression of any given protein for *Vitis amurensis* (red), *V. riparia* (yellow), and *V. vinifera* 'Cabernet Sauvignon' (purple) and 'Riesling' (green) during deacclimation and budbreak. Multiple DEGs encoding for the same protein had their expression aggregated. Arrows indicate transition. For details on graph axes and symbols see Fig. **3**.

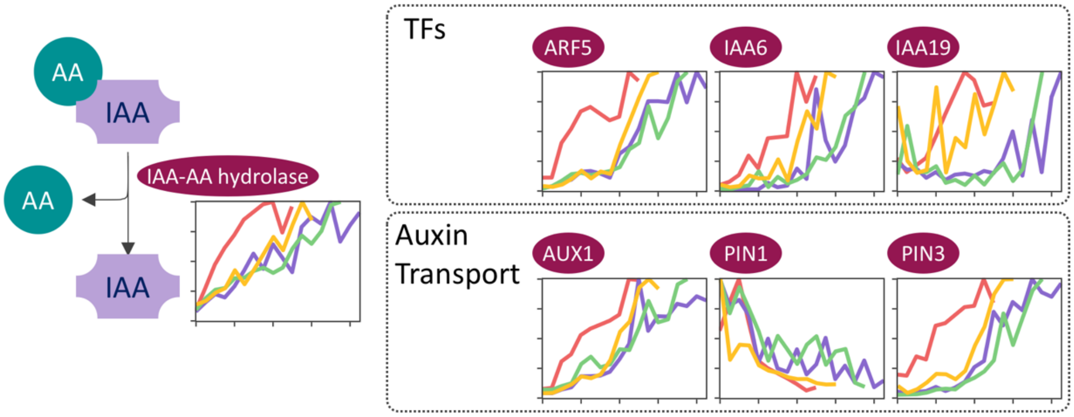

**Fig. S7** Relative expression of genes in the VitisNet circadian rhythm pathway during deacclimation and budbreak in *Vitis amurensis* (red), *V. riparia* (yellow), and *V. vinifera* 'Cabernet Sauvignon' (purple) and 'Riesling' (green).

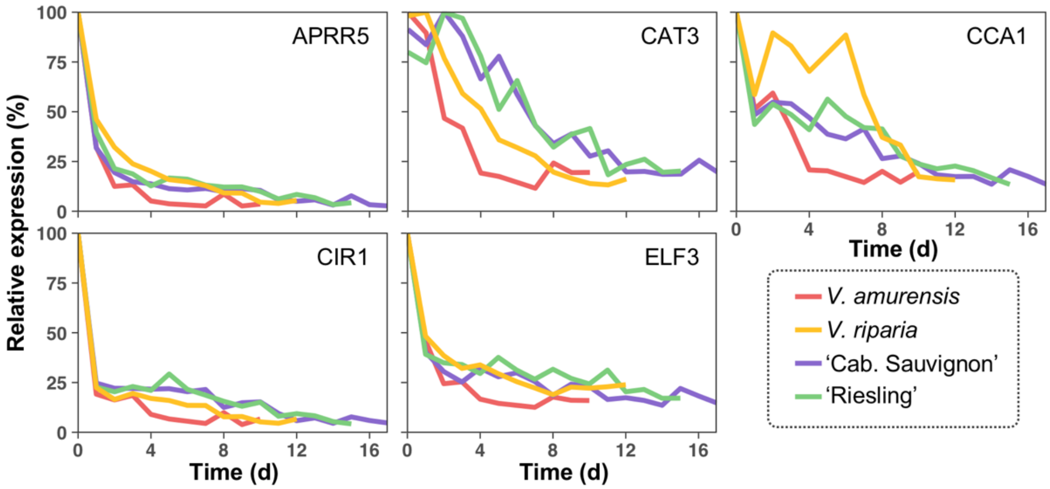

**Fig. S8.** Reduced VitisNet phenylpropanoid biosynthesis pathway. Mini-graphs indicate relative level of expression of any given protein for *Vitis amurensis* (red), *V. riparia* (yellow), and *V. vinifera* 'Cabernet Sauvignon' (purple) and 'Riesling' (green) during deacclimation and budbreak. Multiple DEGs encoding for the same protein had their expression aggregated. Arrows indicate transition. For details on graph axes and symbols see Fig. **3**.

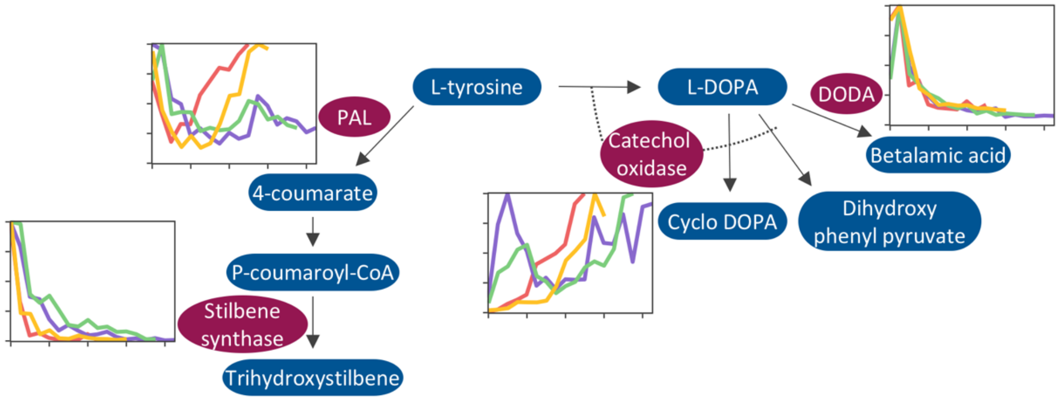

**Fig. S9.** Reduced VitisNet fatty acids synthesis and metabolism pathway. Mini-graphs indicate relative level of expression of any given protein for *Vitis amurensis* (red), *V. riparia* (yellow), and *V. vinifera* 'Cabernet Sauvignon' (purple) and 'Riesling' (green) during deacclimation and budbreak. Multiple DEGs encoding for the same protein had their expression aggregated. Arrows indicate transition. Multiple arrows indicate transition or signaling steps omitted. For details on graph axes and symbols see Fig. **3**.

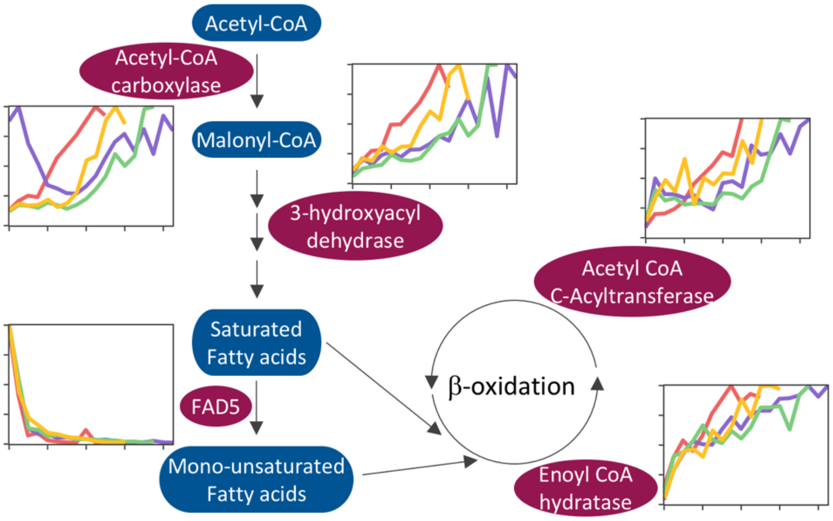

**Fig. S10** Relative expression of genes in the VitisNet photosynthesis antenna proteins and transport electron carriers pathways during deacclimation and budbreak in *Vitis amurensis* (red), *V. riparia* (yellow), and *V. vinifera* 'Cabernet Sauvignon' (purple) and 'Riesling' (green).

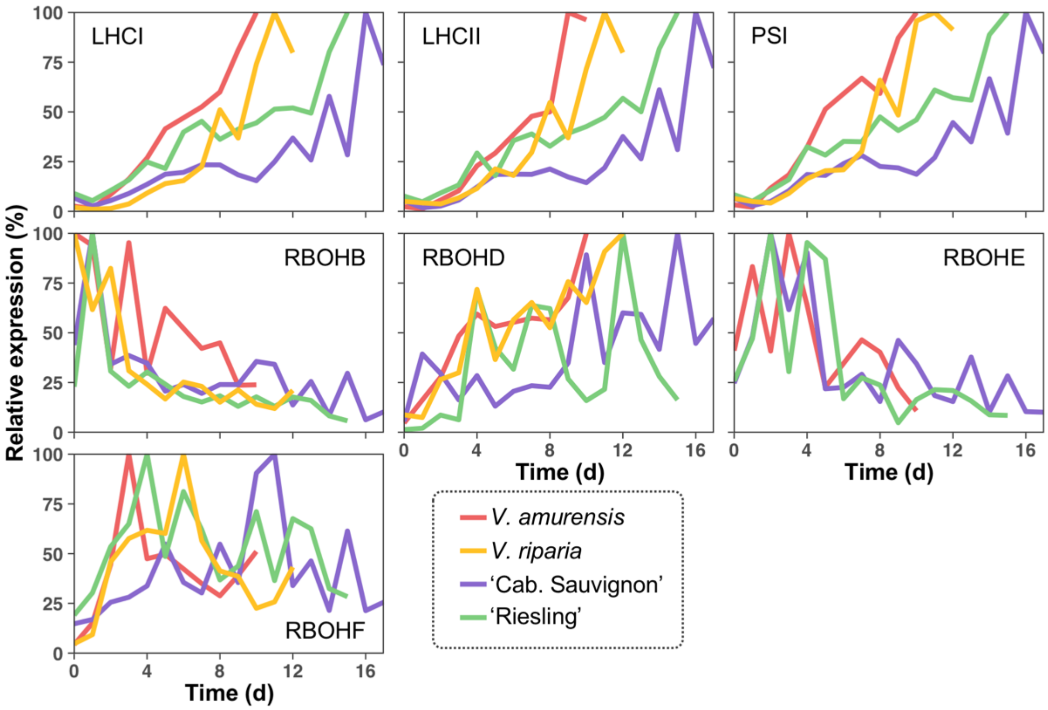
